# Whole slide image analysis of the endometrial decidual reaction reveals multiscale perturbations associated with miscarriage

**DOI:** 10.64898/2026.05.22.727262

**Authors:** George Wright, Thomas M. Rawlings, Mark Eastwood, Paul Brighton, Mireia Taus Nebot, Anja Estermann, William T. M. Flett, Awais Younis, Komal Makwana, Hiroyuki Yoshihara, John D. Aplin, Chow-Seng Kong, Mark Christian, Emma S. Lucas, Joanne Muter, Jan J. Brosens, Fayyaz Minhas

**Affiliations:** Predictive Systems in Biomedicine (PRISM) Lab, University of Warwick, Coventry CV4 7AL, UK; Tissue Image Analytics (TIA) Centre, Department of Computer Science, University of Warwick, Coventry CV4 7AL, UK; Loke Centre for Trophoblast Research, Department of Physiology, Development and Neuroscience, University of Cambridge, Cambridge CB2 3EG, UK; Warwick Medical School, University of Warwick; Coventry CV4 7AL, UK; Department of Biosciences, School of Science and Technology, Nottingham Trent University, Clifton, Nottingham NG11 8NS, UK; The Francis Crick Institute, London NW1 1AT, UK; Division of Developmental Biology & Medicine, School of Medical Sciences, Faculty of Biology, Medicine and Health, University of Manchester, Manchester; M13 9PL, UK; School of Medicine and Population Health, University of Sheffield; S10 2RX, Sheffield, UK; Institute for Precision Diagnostics & Translational Medicine, University Hospitals Coventry & Warwickshire National Health Service Trust; Coventry CV2 2DX, UK

## Abstract

The inflammatory decidual reaction renders the cycling endometrium transiently permissive for embryo implantation before transforming it into the decidua, the maternal bed accommodating the fetal placenta during pregnancy. Disruptions in decidual tissue remodeling are linked to miscarriage and other pregnancy disorders. However, endometrial assessment is hampered by a lack of affordable technologies capable of mapping the spatiotemporal dysregulation of this dynamic and complex tissue. Employing a graph neural network on whole slide images of 493 CD56-immunostained endometrial samples, Endometronome was developed as a deep learning tool to spatially track the decidual reaction and provide accurate estimates of marker gene expression. When applied to 2,690 additional biopsies, this model consistently identified morphological correlates of prior miscarriage burden, a proxy for future risk. Further, a morphological signature indicative of metabolic glandular impairment discriminated between clinical miscarriage presentations. These findings illustrate how advanced imaging analysis of routine histology can transform miscarriage prevention strategies.

## INTRODUCTION

The decidual reaction is a foundational process in mammalian pregnancy, defined as the time-sensitive, progesterone-dependent transformation of the endometrium into the decidua^1–3^. This histotrophic and immunotolerant matrix anchors and supports the invading semi-allogenic placenta throughout gestation. In a few species, including humans and other higher primates, the decidual reaction is not triggered by the implanting embryo but initiated during the midluteal phase of each ovulatory cycle^4–6^. As ovarian progesterone production decreases rapidly in the absence of implantation, decidual transformation of the endometrium is abandoned during the late luteal phase of non-conception cycles, resulting in an influx of macrophages and granulocytes, degradation of the superficial layer, and menstrual bleeding^7,8^.

The human endometrium comprises about 1000 mm^2^ of luminal epithelium. This epithelium connects to roughly 15,000 glands within a vascularized stroma rich in immune cells^9^. Following menstrual repair, rising ovarian and local estradiol production typically quadruples endometrial volume and restores the functional layer prior to ovulation^10^. Local morphogens and cytokines form gradients that drive angiogenesis and regulate cell proliferation, thereby establishing positional cell identities and regional patterns^3,11,12^. After ovulation, progesterone inhibits proliferation, drives metabolic reprogramming, and promotes endometrial cell differentiation^7^. These processes result in a sudden change in endometrial gene expression five days later^13^, marking the start of the inflammatory decidual reaction and the opening of a four-day implantation window^3,7,11^. Histologically, the decidual reaction is defined by the onset of apocrine glandular secretions, localized edema, and proliferation and differentiation of CD56+ uterine natural killer (uNK) cells into immunotolerant subsets^14,15^. In parallel, the progressive loss of subluminal implantation niche cells, concurrent with the emergence of decidual cells, leads to the closure of the window^11^. Decidual cells, which exhibit epithelioid morphology and a stress-resistant phenotype, arise from transcriptionally reprogrammed, progesterone-dependent stromal fibroblasts^16^.

Upon embedding of the blastocyst in the subluminal implantation niche^17^, emerging placental trophoblast cells not only invade the decidualizing stroma and glands but also the terminal spiral arterioles^18,19^, thereby creating a protective low-oxygen environment at the nascent maternal-fetal interface. Consequently, rapid placental growth and fetal organogenesis depend on an abundance of glandular secretions rich in glucose, lipids, glycoproteins, and growth factors^20^. By the end of the first trimester, the loss of vascular trophoblast plugs leads to placental perfusion and a dramatic rise in oxygen levels^21^. Continued intravascular trophoblast invasion transforms the uterine spiral arteries into large fibrinoid vessels, ensuring that placental perfusion meets the metabolic demands of the developing fetus^22,23^.

Compelling evidence indicates that dysregulation of the decidual reaction, including acceleration and stalling, leads to reproductive failure^11,24,25^. Miscarriage, defined as the loss of a pregnancy before viability, occurs in over 90% of cases during the first trimester^26^. The estimated pooled risk of miscarriage in all clinically recognized pregnancies is approximately 15%^27^. Maternal age is a key determinant of baseline risk, reflecting the increased incidence of meiotic chromosome errors in oocytes and embryos among women over 35 years. Each pregnancy loss further increases the risk by 5-10%^26,28,29^. Recent findings indicate that the recurrence risk associated with prior losses closely aligns with the frequency of decidual reaction stalling across menstrual cycles^11^. Lack of decidual propagation extends the implantation window, disrupts embryo-endometrium synchrony, and results in an unstable placental-decidual interface, which can lead to catastrophic bleeding. Perturbations in the cyclical decidual reaction have also been observed following preeclamptic pregnancies^30,31^. These observations suggest that menstrual cycles form recursive loops, in which detrimental outcomes from a previous cycle, such as miscarriage and obstetrical disorders, can adversely impact cellular dynamics in subsequent cycles, thereby increasing the risk of further adverse pregnancy outcomes^11^.

Since the decidual reaction is initiated during each ovulatory cycle, it may be possible to identify individuals at risk of adverse pregnancy outcomes prior to conception and to develop pre-pregnancy therapeutics. This strategy could mitigate, at least partly, the lack of pharmacological innovation in maternal medicine^32^, a legacy of fetal harm caused by medication taken in pregnancy, such as thalidomide, sodium valproate, and diethylstilbestrol^33–35^. However, the development of pre-pregnancy endometrial diagnostics faces several challenges, including the requirement for large sample sets to define the physiological boundaries of the inflammatory decidual reaction and the need for cost-effective technologies that facilitate comprehensive spatiotemporal analysis of this dynamic process in clinical settings. Among potential approaches, histology whole slide images (WSIs) of tissue samples capture rich spatial and morphological information that reflects underlying gene regulatory states^36,37^. Recent advances in machine learning have transformed the analysis of complex histological data, enabling the identification of clinically relevant patterns that were previously inaccessible^38–40^. To address the lack of pre-conception endometrial diagnostics, we developed Endometronome, a deep learning tool trained on 493 CD56-immunostained endometrial samples, validated in 160 independent samples, and applied to 2,690 additional biopsies, capturing a total of 1.19 billion cells, 2.64 million gland components, and 9.3 million image patches for analysis. Our findings demonstrate that Endometronome is a powerful automated tool for comprehensive, multiscale profiling of the pre-pregnancy decidual reaction.

## RESULTS

### Multiscale graph analysis of endometrial WSIs

Endometrial pipelle biopsies, sampling the upper endometrial layer at 1–2 mm depth, were obtained 5–12 days after the pre-ovulatory luteinizing hormone surge (LH+5 to +12), as determined by home ovulation testing (Figure 1A). Formalin-fixed paraffin-embedded (FFPE) samples were processed for CD56 (neural cell adhesion molecule 1) immunohistochemistry with 3,3′-diaminobenzidine (DAB) as the chromogen and hematoxylin counterstain. In parallel, transcript levels of 13 genes. enriched in epithelial, stromal, or uNK cells (Figure S1), were measured by RT-qPCR in 493 samples (Figure 1B).

**Figure 1.**
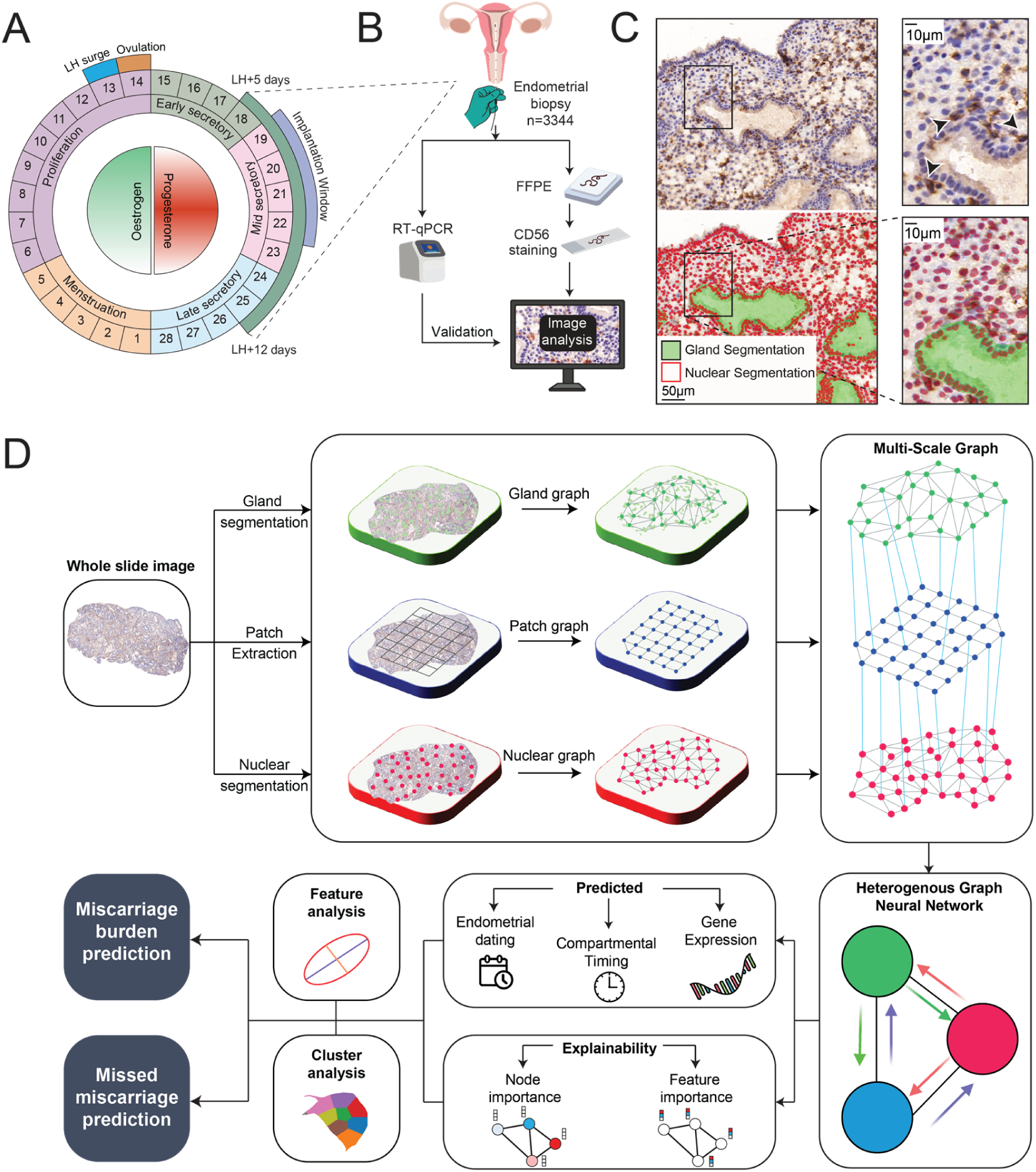
Endometronome framework for multiscale analysis of the decidual reaction from whole-slide histology. (A) Schematic overview of the menstrual cycle illustrating hormonal dynamics, luteal-phase progression, and the timing of endometrial biopsies relative to the luteinizing hormone (LH) surge. Samples spanning the proliferative phase through the mid- and late-secretory phases were analysed. (B) Experimental workflow. Endometrial biopsies were processed as formalin-fixed paraffin-embedded (FFPE) tissue sections for CD56 immunohistochemistry and whole-slide imaging, alongside matched RT-qPCR profiling for molecular timing and gene expression analyses. (C) Representative whole-slide image regions showing automated segmentation of glandular structures (green) and nuclei (red). Insets illustrate the detection of epithelial gland boundaries and nuclear morphology used for downstream feature extraction. (D) Overview of the Endometronome computational pipeline. Whole-slide images were decomposed into multiscale representations comprising gland, patch, and nuclear graphs. These subgraphs were integrated into a unified heterogeneous graph neural network (HGNN) framework enabling information exchange across spatial scales. The HGNN was trained to predict endometrial timing, compartment-specific decidual progression, and gene expression directly from histology while simultaneously supporting interpretable spatial analysis through node- and feature-level attribution. Downstream analyses linked inferred tissue states to miscarriage burden and recurrent missed miscarriage-associated endometrial phenotypes.

Using these 493 WSIs with matched gene expression, we developed Endometronome, a machine learning framework that segments individual nuclei, glands, and larger tissue patches (Figure 1C), extracting key features such as nuclear morphology, glandular organization, and tissue architecture illustrated in the pictorial glossary in Figure S4. These multiscale descriptors are integrated into a heterogeneous graph representation of the tissue, enabling the model to capture spatial relationships across multiple scales (Figure 1D and Figure S2). The resulting graph is then used to predict endometrial features, including sample dating, progression of the decidual reaction in different cellular compartments, and gene expression. Model performance was assessed using internal five-fold cross-validation and tested on an independent set of 160 samples (Figure S3).

We subsequently applied Endometronome to an additional 2,945 biopsies to investigate morphological correlates of miscarriage burden and clinical presentation (Figure 1D and Figure S3). Across all cohorts, the analysis encompassed 1.19 billion cells, 2.64 million glandular structures, and 9.3 million image patches. Demographic characteristics of study participants are summarized in Table S1.

Importantly, Endometronome incorporates feature-attribution analysis using GNNExplainer to identify the image regions and histological structures driving model predictions, enabling interpretable spatial mapping of inferred tissue states (Figure S2). Combined with differential abundance and enrichment analyses, this framework enabled the discovery of statistically significant glandular and nuclear states associated with decidual progression, miscarriage burden, and recurrent missed miscarriage. Interactive visualization of Endometronome spatial predictions across WSIs is available at https://endometronome.dcs.warwick.ac.uk/.

### Endometrial dating, compartmental timing, and gene expression predictions

Histological dating of the endometrium traditionally relies on the Noyes criteria, a set of morphological features established over 75 years ago^14^. To determine whether Endometronome could recover temporal progression of the decidual reaction directly from histology, we predicted both patient-reported cycle timing (LH+) and EndoTime, a transcriptomic framework for continuous luteal-phase dating based on six genes (*IL2RB*, *IGFBP1*, *CXCL14*, *DPP4*, *GPX3*, and *SLC15A2*)^41^. Endometronome predictions showed strong concordance with measured timing during both cross-validation and independent testing, with EndoTime predictions achieving Spearman’s ρ = 0.78 ± 0.04 and 0.74 ± 0.05, respectively, outperforming ovulation test-based dating (Figure 2A and Table S2).

**Figure 2.**
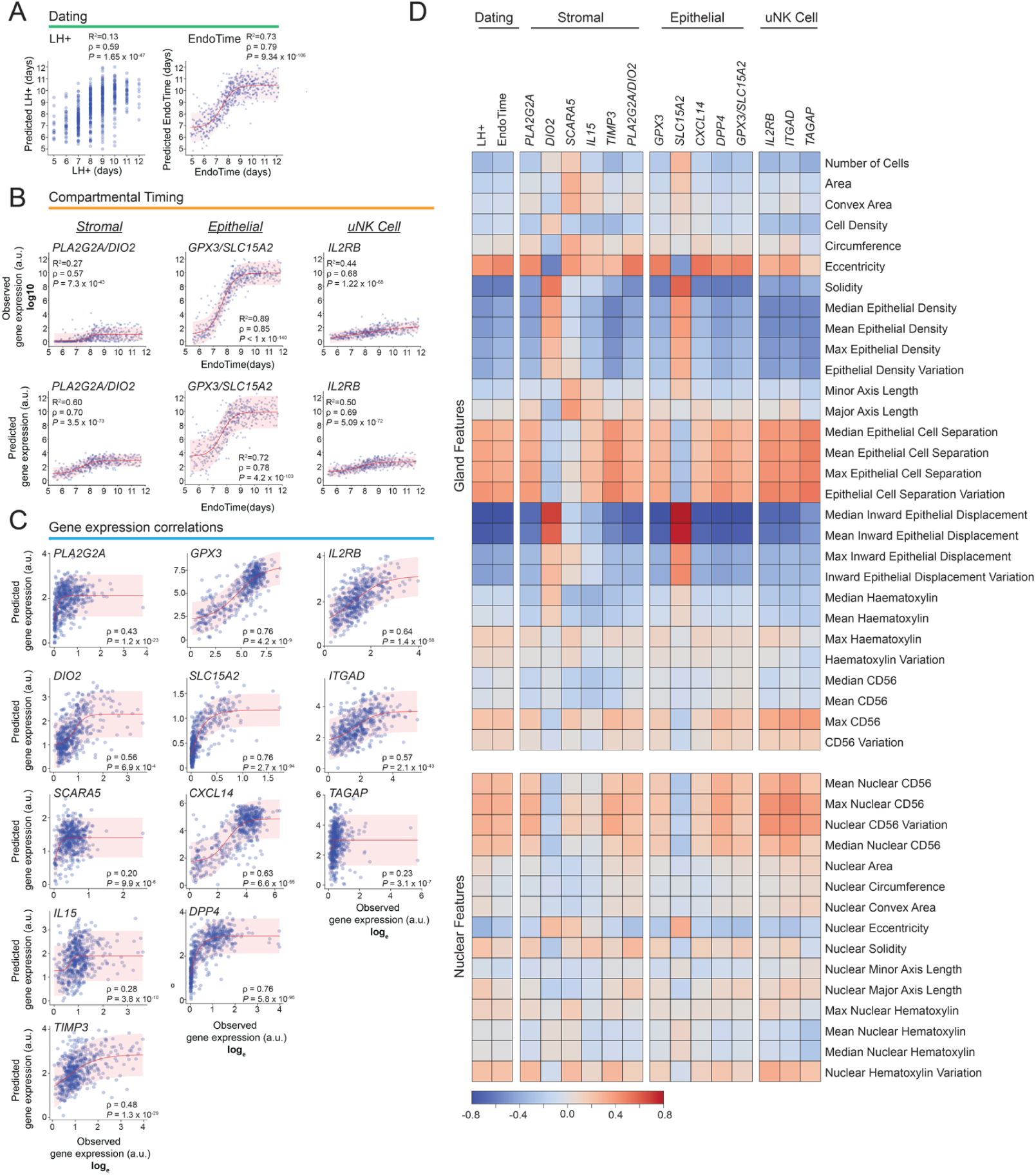
Endometronome predicts endometrial dating, compartmenttiming, and gene expression directly from whole-slide histology. (A) Predicted versus observed endometrial timing for patient-reported luteinizing hormone day (LH+) and EndoTime transcriptomic dating. Each point represents an individual biopsy sample. Solid lines indicate fitted regression curves with shaded 90% confidence intervals. Spearman’s rank correlations and associated p values are shown for each prediction task. (B) Predicted versus observed compartment-specific timing measures reflecting stromal (*PLA2G2A/DIO2*), epithelial (*GPX3/SLC15A2*), and uterine natural killer (uNK) cell (*IL2RB*) progression during the decidual reaction in cross-validation and independent test cohorts. Endometronome accurately recapitulates molecular signatures of coordinated decidual remodeling across cellular compartments; a.u., arbitrary units. (C) Predicted versus measured expression values for representative genes associated with stromal, epithelial, and immune decidual programs. Glandular epithelial genes, including *GPX3*, *CXCL14*, *DPP4*, and *SLC15A2*, exhibited the strongest concordance between histology-derived predictions and RT-qPCR measurements. (D) Correlation heatmap linking interpretable glandular and nuclear morphological features with predicted timing and gene expression targets. Rows correspond to histological features grouped by similarity, and columns correspond to molecular and temporal targets grouped by functional relationship. Red and blue indicate positive and negative correlations, respectively. The analysis reveals coordinated associations between glandular architecture, epithelial organization, nuclear morphology, and progression of the decidual reaction across stromal, epithelial, and immune compartments.

Histological dating assigns each sample a cycle day, which can mask endometrial asynchrony, defined as the misalignment of glandular maturation, stromal remodeling, and uNK cell expansion and differentiation^11,42^. To resolve these compartment-specific dynamics, molecular timing approaches have been developed for epithelial, stromal, and immune compartments. For example, stromal remodeling can be evaluated using the *PLA2G2A*-to-*DIO2* transcript ratio^11^. When normalized to the cycle day of endometrial sampling, this ratio tracks the time-dependent loss of progesterone-resistant *DIO2*+ implantation niche cells and progesterone-responsive *PLA2G2A*+ predecidual cells expand (Figure 2B). In the glands, the ratio of *GPX3* to *SLC15A2* transcripts increases exponentially as the endometrium progresses through the implantation window, reflecting reciprocal gene regulation^16^. *IL2RB* expression is a putative marker of the time-dependent expansion and differentiation of uNK cells (Figure S1). We next asked whether these molecular measures of compartmental timing are encoded in endometrial morphology and can be recovered from WSIs using Endometronome.

Endometronome predictions showed strong concordance with molecular markers of decidual progression across all three compartments (Figure 2B and Table S2). Predictive performance was highest for glandular markers, including *SLC15A2*, *DPP4*, *GPX3*, and *CXCL14* (Figure 2C and Table S2), suggesting that glandular morphology undergoes particularly coordinated structural and transcriptional remodeling during decidualization. Since each endometrial gland originates from a single progenitor cell^43^, the effects of the inflammatory decidual reaction on glandular morphology and gene expression are likely to be more spatially coordinated and transcriptionally coherent than the corresponding changes in stromal or immune compartments (Figure 2B). Strong concordance was also observed for *ITGAD*, a marker of immunotolerant uNK subsets^11^, and for *PLA2G2A* and *TIMP3* expression in decidualizing stromal cells. By contrast, *TAGAP*, *SCARA5*, and *IL15* demonstrated moderate performance.

To evaluate the contribution of multiscale integration to these predictions, Endometronome was systematically compared with its constituent patch-, cell-, and gland-level graph architectures. Across the 16 biomarker ratio and gene expression targets, the integrated heterogeneous graph model achieved the highest predictive accuracy for 11 variables, while performance differences for the remaining targets were minimal (Table S2). Analysis of individual subgraph models revealed scale-specific predictive specialization, with patch- and gland-level graphs each achieving the strongest performance for seven variables, whereas cell-level graphs performed best for two targets. These findings suggest that distinct histological scales capture complementary aspects of decidual remodeling and that their integration within the HGNN provides a more comprehensive representation of endometrial tissue dynamics.

Analysis of the morphological features driving Endometronome predictions revealed a continuous trajectory of endometrial differentiation, linking glandular architecture to compartment-specific decidual programs (Figure 2D and Figure S2B). Key glandular features, including solidity, eccentricity, epithelial cohesion, and epithelial positioning relative to the gland boundary, are illustrated in the pictorial glossary in Figure S4. Feature-attribution analysis identified that these glandular and nuclear characteristics as major contributors to model predictions. During the decidual reaction, high prediction scores were associated with progressive gland elongation and irregularity, characterized by reduced solidity, increased eccentricity, decreased epithelial cohesion, and more superficial epithelial positioning. These changes, consistent with partial pseudo-vacuolization of glandular epithelium and luminal expansion, coincided with local stromal and immune activation, reflected by increased CD56 immunoreactivity. By contrast, pre-decidual states associated with high *DIO2* and *SLC15A2* expression exhibited round, cohesive glands with dense epithelial packing, narrow lumina, and minimal pseudo-vacuolization. At the nuclear level, increased nuclear size and CD56 immunoreactivity were also strongly associated with decidual progression (Figure 2D). Together, these findings indicate that Endometronome captures coordinated morphological transitions underlying dynamic decidual remodeling of the endometrium.

### Spatial heterogeneity and clustering of glandular and nuclear states

In contrast to conventional gene expression analysis, which produces a single composite value per sample, EndoMetronome enables spatial mapping of spatially localized node-level predictions on WSIs. Spatial heterogeneity, depicted as heatmaps with scores ranging from low (blue) to high (red), was evident across dating, compartmental timing, and gene expression predictions (Figure 3 and Figure S5), highlighting the influence of local cues in driving decidualization. As shown in Figure 3, divergence in compartmental timing, a measure of decidual progression in distinct cell types, becomes more prominent upon closure of the putative implantation window. Figure S5 presents spatial maps of predicted gene expression. As expected, induction of glandular genes (*GPX3*, *DPP4,* and *CXCL14*) was more homogeneous and prominent than gene changes associated with stromal or uNK cell differentiation. Gene expression predictions that correlated only modestly with measured transcript levels, such as *IL15*, *SCARA5*, and *TAGAP* (Figure 2C), show prominent regional variation at the start of the implantation window (LH+5/6 days) and less clear temporal patterns during the decidual reaction.

**Figure 3.**
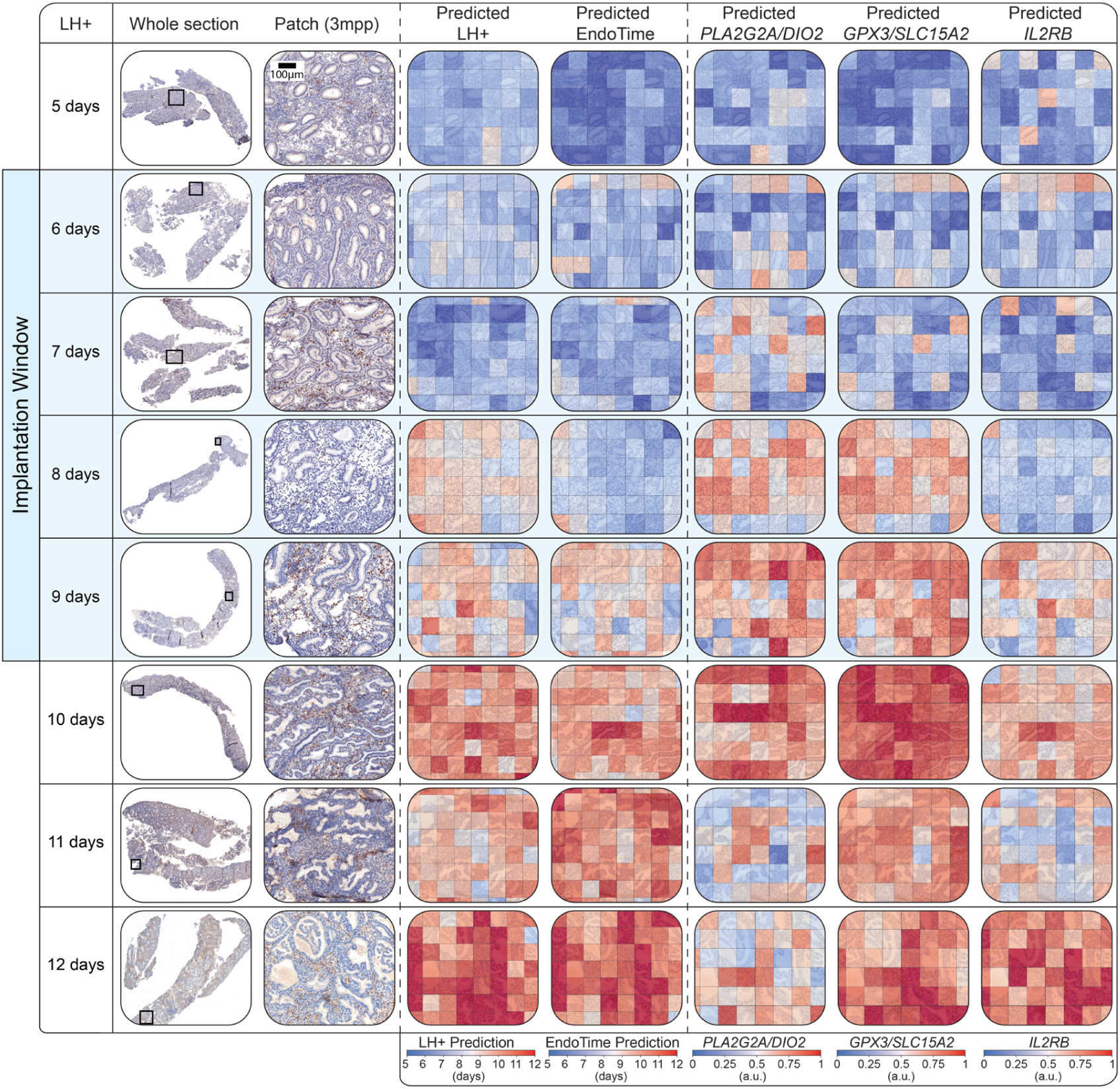
Spatial prediction maps reveal progressive and heterogeneous decidual remodeling across the luteal phase. Representative whole-slide image regions and corresponding Endometronome prediction maps across the implantation window. Rows correspond to biopsies obtained at different days following the luteinizing hormone (LH) surge, illustrating temporal progression of the decidual reaction. For each sample, the left panels show the whole-slide section and a representative high-resolution tissue patch at 3 microns per pixel (mpp) including a 100μm scale bar. Spatial prediction maps display localized patch-level estimates for LH+, EndoTime, stromal timing (*PLA2G2A/DIO2*), epithelial timing (*GPX3/SLC15A2*), and uNK cell timing (*IL2RB*). Blue and red indicate low and high predicted values, respectively. Endometronome captures coordinated but spatially heterogeneous progression of decidual transformation across stromal, epithelial, and immune compartments, with increasingly elevated and spatially structured predictions observed during progression through the implantation window.

To determine whether these spatially heterogeneous predictions reflected recurrent morphological states during decidualization, glandular and nuclear architectures were clustered across WSIs, yielding ten distinct glandular states (Figure 4A, 4B, and Figure S6). The relative abundance of each cluster within individual samples was then analyzed against endometrial timing, enabling identification of glandular states significantly enriched or depleted across progression of the decidual reaction. Early-luteal glandular states were characterized by compact glands with high solidity and dense epithelium, whereas later states exhibited progressive gland elongation, epithelial separation, and luminal expansion (Figure 4C). Consistent with progression through the decidual reaction, clusters 1 to 4 progressively declined across the luteal phase, while clusters 6 to 9 became increasingly abundant. These coordinated transitions closely mirror the classical morphological progression underlying Noyes endometrial dating criteria^14^ and suggest that Endometronome captures recurrent glandular remodeling states associated with decidual transformation.

**Figure 4.**
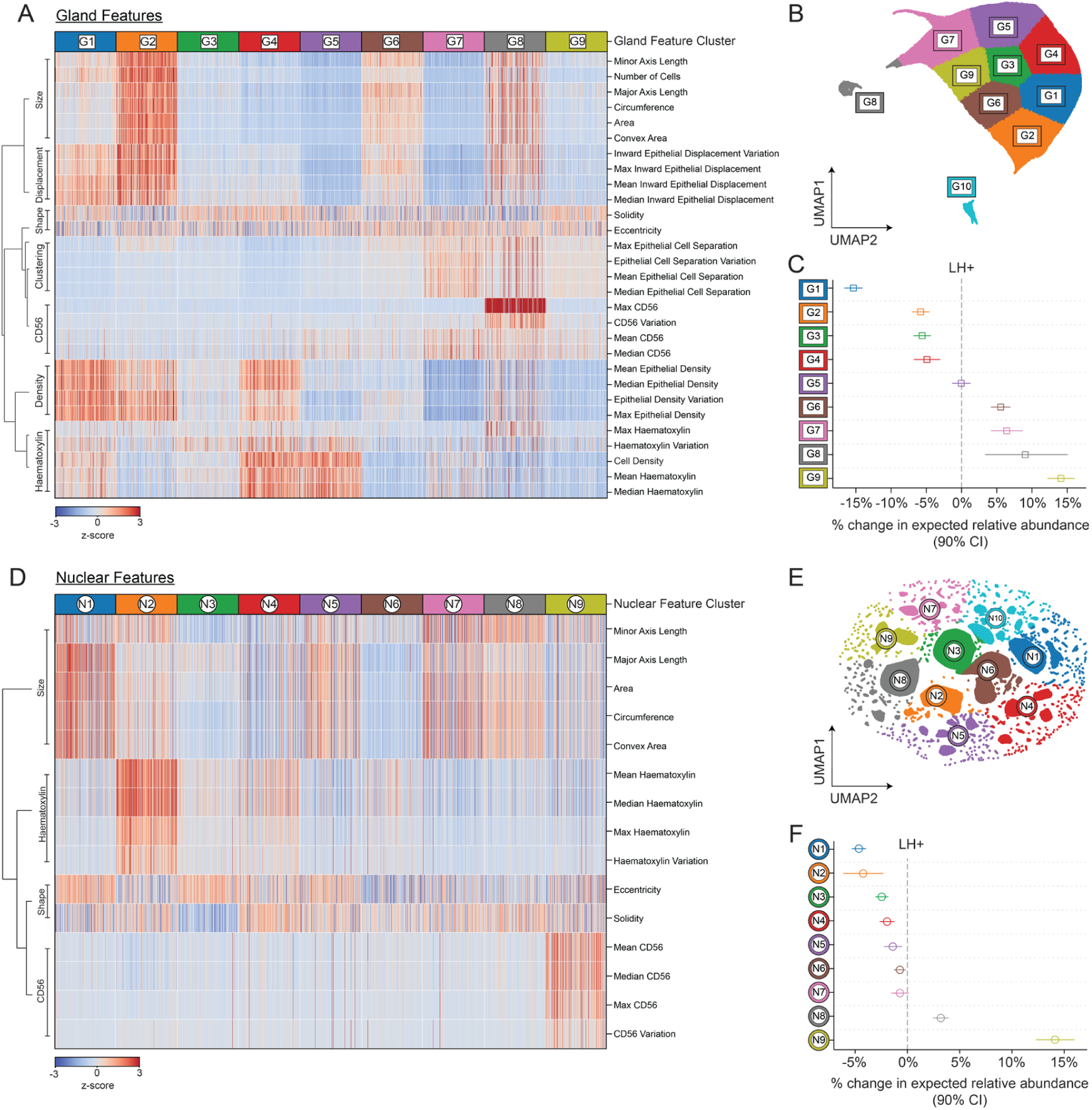
Distinct glandular and nuclear tissue states exhibit coordinated temporal remodeling during the decidual reaction. (A) Heatmap of z score-normalized glandular feature profiles across identified gland communities (G1–G9). Rows correspond to interpretable glandular features hierarchically clustered by similarity, and columns represent individual gland instances grouped by k-means community assignment. Communities are ordered according to their association with progression through the luteal phase. Features linked to glandular maturation, including gland eccentricity, epithelial separation, and lumen expansion, progressively increase across temporally enriched gland states, whereas compact glandular morphologies characterized by high solidity and epithelial density predominate in early-phase communities. (B) UMAP visualization of gland clusters revealing distinct gland communities. Spatial separation of clusters indicates recurrent glandular morphologies associated with different stages of decidual remodeling. (C) Differential abundance analysis using regression of cluster proportions against time demonstrates time-dependent compositional changes, with cluster 9 increasing by approximately 15% per unit time across the luteal phase showing the temporal changes in gland clusters. (D) Heatmap of z-score–normalized nuclear features, organized as in (A). Cluster N9 shows high CD56 staining associated with uNK cell presence. (E) UMAP visualization of nuclear clusters showing distinct associations with temporal changes. (F) Differential abundance analysis using regression of nuclear cluster proportions against time illustrating temporal variation in nuclei clusters with cluster 9 increasing by 15% per unit time.

Clustering of nuclear morphology revealed substantially greater diversity, reflecting the complex and dynamic cellular composition of the cycling endometrium (Figure 4D and 4E). Analysis of nuclear cluster abundance across timed biopsies identified multiple nuclear states significantly associated with progression through the decidual reaction (Figure 4F). Clusters 1 to 5, composed predominantly of elongated nuclei, progressively declined across the luteal phase, indicating reduced prevalence of the corresponding cellular states during decidual transformation. By contrast, clusters 8 and 9 became significantly enriched over time, with Cluster 9 exhibiting strong CD56 immunoreactivity, consistent with continued expansion of uNK cells in late-luteal endometrium prior to menstruation (Figure 4D).

### Persistent glandular and nuclear tissue states associated with miscarriage burden

Reproductive success is determined not only by embryo fitness, which declines with advancing maternal age^28^, but also on the sustained calibration of the decidual reaction across conception cycles^11^. Recent evidence suggests that loss of uterine plasticity due to prolonged tissue inflammation associated with miscarriages impairs the calibration and propagation of the decidual reaction in subsequent cycles^11,44^. To investigate whether these persistent perturbations are encoded in endometrial morphology, Endometronome was applied to an additional 2,690 biopsies, establishing a cohort of 3,226 timed endometrial samples (Figure S3).

Endometronome predictions and histological features were normalised for the day of endometrial sampling by establishing percentile ranks within a fitted normal distribution. Normalised predicted *PLA2G2A*/*DIO2* ratios, which track the decidual reaction in stromal cells, demonstrated a strong inverse association with miscarriage burden, defined as the number of prior losses (Figure 5A). Consistent with epidemiological data^26,28^ and previous molecular studies^11^, increasing miscarriage burden was associated with a stepwise decline in both predicted and observed scaled *PLA2G2A*/*DIO2* ratios, indicative of impaired decidual propagation. Univariate Bayesian modeling demonstrated that higher predicted *PLA2G2A/DIO2* ratios were consistently associated with lower miscarriage burden, with each 10-percentage-point increase corresponding to a 2.3% reduction in expected miscarriage incidence (incidence rate ratio (IRR) = 0.976, 90% credible interval (CrI) = 0.974–0.993), replicating the ground-truth ratio effect (IRR= 0.947, 90% CrI = 0.922–0.973). Modeling of measured gene expression and Endometronome-derived predictions revealed coordinated impairment of glandular, stromal, and immune decidual programs with increasing miscarriage burden, reflected by reductions in normalized *GPX3/SLC15A2* ratios and *IL2RB* expression (Figure 5A). All genes demonstrated measurable effects: early-luteal genes, including *SLC15A2*, were positively associated with miscarriage burden (median IRR = 1.017, 90% CrI = 1.007–1.027), whereas *SCARA5*, a highly progesterone-responsive stromal gene^45^, showed a strong protective association (median IRR = 0.974, 90% CrI = 0.968–0.983). Endometronome-derived predictions yielded narrower credible intervals than transcript-based analyses, likely reflecting the substantially larger histological cohort size (Figure 5A).

**Figure 5.**
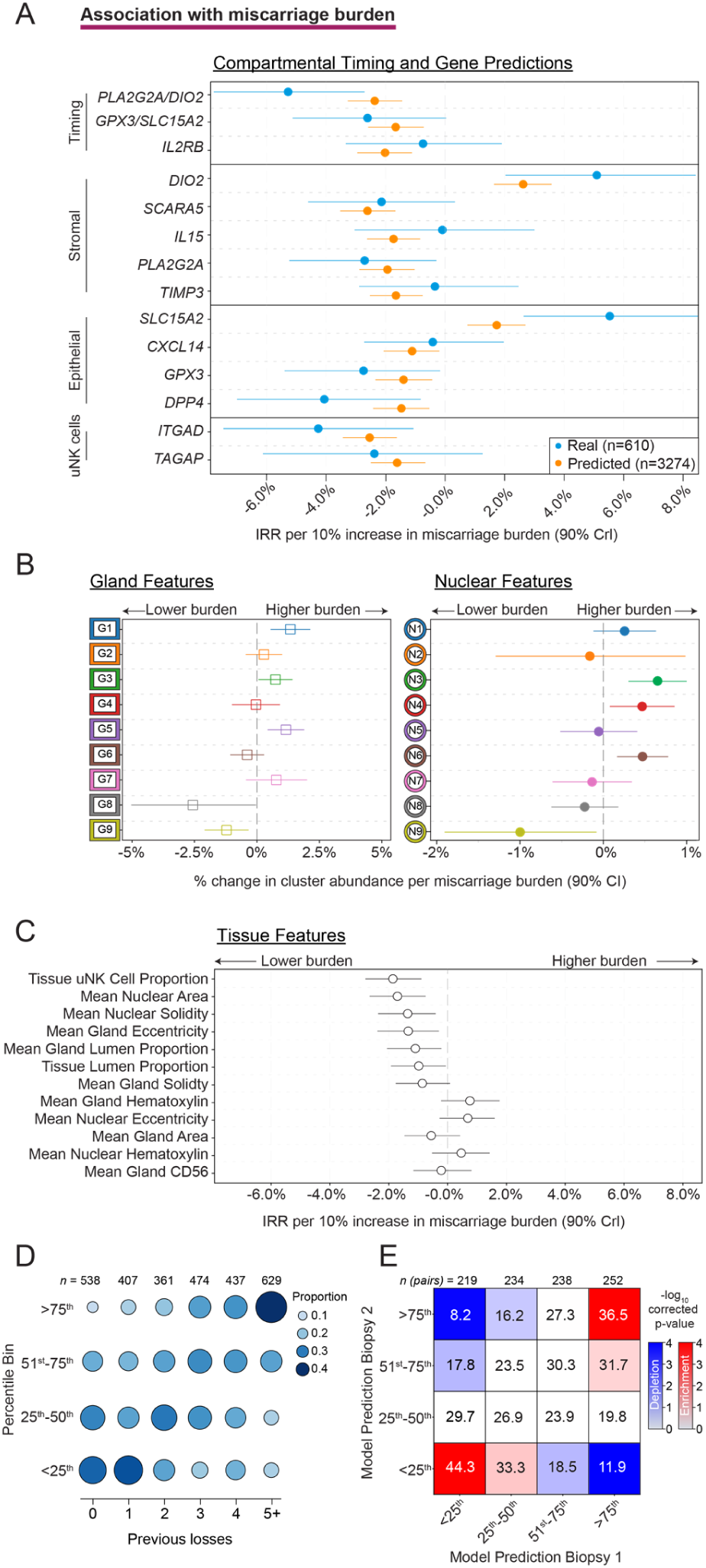
Persistent glandular and nuclear tissue alterations associated with miscarriage burden. (A) Bayesian linear regression coefficients (90% credible intervals) for ground-truth and predicted gene expression with miscarriage burden. Showing the *PLA2G2A/DIO2* ratio decreasing by around 4% with each 10% increase in miscarriage burden. (B) Differential community abundance analysis illustrates significant differences in glandular and nuclear cluster composition with respect to miscarriage burden (90% confidence interval). (C) Bayesian linear regression coefficients (90% credible intervals) for interpretable image features with miscarriage burden reinforcing that there are distinct differences in feature presentation following increasing miscarriages. (D) Dot plot showing the percentage of model predictions falling into each bin for different miscarriage burdens showing the model predictions falling within the >75th percentile bin increase with each subsequent miscarriage. (E) Contingency tables show recurrence rates of predicted model score in paired biopsies. The coloured squares in the contingency tables indicate statistical significance (P < 0.05), as determined by Fisher’s exact test for enriched (red key) and depleted (blue key) associations showing the recurrence of the model predictions across cycles.

We next examined whether miscarriage burden was associated with specific glandular and nuclear tissue morphological states. After adjusting for endometrial timing, multiple glandular clusters exhibited significant abundance shifts with increasing miscarriage burden. Clusters 8 and 9, associated with well-organized glandular architecture and intact epithelial-stromal interfaces, were progressively depleted with increasing miscarriage burden, suggesting a protective association (Figure 5B, left panel, and S7A). By contrast, clusters 1, 3, and 5 displayed the opposite pattern, with greater abundance associated with higher miscarriage risk. Clusters 1 and 3 were characterized by morphologically stunted glands with high solidity and low eccentricity, features consistent with restricted or delayed decidual progression. These findings support the concept that the persistence of morphological features characteristic of the early luteal phase endometrium reflects blunting or stalling of the decidual reaction. Notably, cluster 5 showed strong association with miscarriage burden despite lacking temporal variation across the luteal phase (Figure 4C), suggesting that it may represent a time-independent morphological biomarker of recurrent miscarriage risk (Figure 4C and 5B, left panel).

Nuclear tissue states also shifted systematically with miscarriage burden. Reduced abundance of nuclear cluster 9, characterized by strong CD56 immunoreactivity (Figure 4D and S7B), was associated with increasing prior losses, consistent with impaired expansion of uNK cells during decidualization (Figure 5B, right panel and S7B).

We next examined whole-tissue morphological features associated with miscarriage burden (Figure 5C). After normalization for endometrial timing, greater gland eccentricity and higher overall glandular tissue proportion, features indicative of advanced glandular maturation and preserved structural organization, were associated with fewer prior losses. Increased mean nuclear size, a canonical hallmark of decidualization^46^, and higher abundance of uNK cells was likewise associated with a lower miscarriage burden (Figure 5C).

To determine whether endometrial morphology encodes a quantifiable signature of miscarriage risk imparted by prior pregnancy losses, we trained a multivariate Bayesian generalized linear model, integrating day-normalized image features and Endometronome-derived predicted timing and gene expression scores. The proportion of samples with predicted high-risk scores (>75^th^ percentile) increased monotonically with the number of prior miscarriages, while representation in the low-risk score category (<25^th^ percentile) declined accordingly (Figure 5D).

Analysis of paired biopsies obtained across independent menstrual cycles from 943 women further demonstrated high recurrence of both high-risk and low-risk tissue states (Figure 5E). Among biopsies assigned high-risk scores (>75^th^ percentile), 36.5% of paired samples from the same individuals were also classified as high risk, significantly exceeding random expectation (Bonferroni-adjusted *p* < 2.48 × 10^−9^, Fisher’s exact test). Similarly, 44.3% of low-risk biopsies (<25^th^ percentile) remained low risk across independent cycles (Bonferroni-adjusted *p* < 4.78 × 10^-11^, Fisher’s exact test). Comparable persistence patterns were observed for selected glandular and nuclear morphological features (Figure S8). Collectively, these findings indicate that prior miscarriages disrupt propagation of the decidual reaction across cycles, leaving persistent perturbations detectable directly from routine histology at the levels of tissue architecture, cellular composition, and inferred molecular state.

### Distinct glandular and nuclear histomorphological features associated with recurrent missed miscarriage

Miscarriage encompasses a heterogeneous spectrum of clinical presentations^27^, which often but not always vary between pregnancy losses. Recurrent patterns can be broadly divided into losses preceded by prominent vaginal bleeding, indicative of breakdown of the placental-decidual interface, and those that are not. Although detailed clinical phenotyping was unavailable for most biopsies, our cohort included 104 women with recurrent missed miscarriage (RMM) (Figure S3), defined here as repeated ultrasound-based detection of early fetal demise in three or more pregnancies in otherwise asymptomatic women (Table S1).

Analysis of glandular and nuclear tissue states revealed that RMM is associated with morphological perturbations distinct from those linked to overall miscarriage burden. For instance, increased prominence of gland cluster 3 and nuclear cluster 3, normalized for cycle day, was associated with a higher miscarriage burden (Figure 5B, S7C, and S7D) but with a reduced likelihood of RMM (Figure 6A). Conversely, gland cluster 6 demonstrated strong enrichment in RMM despite lacking association with general miscarriage burden, suggesting that RMM is associated with a distinct endometrial tissue state.

**Figure 6.**
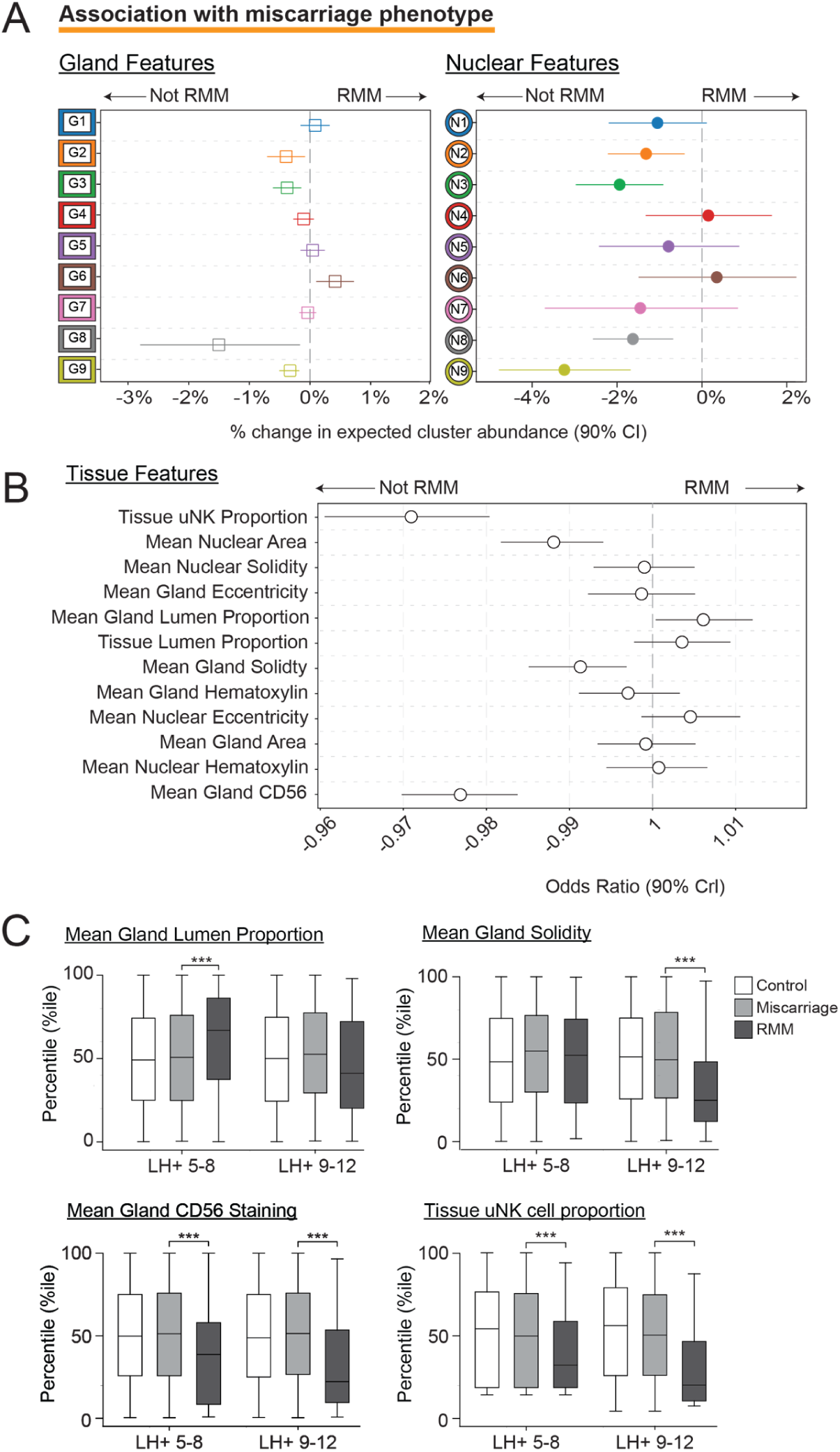
Distinct glandular and immune tissue states associated with recurrent missed miscarriage (RMM). (A) Differential community abundance analysis illustrates significant differences in glandular and nuclear cluster composition with respect to RMM (90% confidence interval). (B) Bayesian linear regression coefficients (90% credible intervals) for interpretable image features with RMM reinforcing that there are distinct differences in feature presentation. (C) Differences in feature distributions across control, miscarriage, and RMM groups. There is a temporal pattern with changes in feature value associated with early and late timing for gland lumen proportion and solidity respectively, whereas gland CD56 staining and tissue uNK cell proportions show alterations in RMM patients throughout the luteal phase.

To further investigate potential histological markers of RMM, Bayesian logistic regression analyses were conducted, focusing on glandular and cellular features (Figure 6B). The proportion of tissue uNK cells exhibited the strongest protective effect for RMM (OR = 0.971; 90% CrI 0.961-0.980), with mean gland CD56 immunoreactivity also ranking high. High normalized mean gland solidity was protective against RMM (OR: 0.991, 90% CrI 0.985-0.997), whereas a reversed association was observed for mean gland lumen proportion (OR: 1.006, 90% CrI 1.001-1.012) (Figure 6B). This indicates impaired glandular cohesion and altered epithelial remodeling in affected individuals.

To determine whether these glandular and immune abnormalities exhibited distinct temporal dynamics during decidualization, we quantified normalized tissue-level features across the early and late phases of the decidual reaction in control subjects without prior losses (n = 607), non-phenotyped miscarriage cases (n = 2,515), and women with recurrent missed miscarriage (RMM) (Table S1 and Figure 6C). A temporal pattern was observed, with enhanced mean gland lumen proportion and lower mean gland solidity in RMM confined to the early and late phases of the decidual reaction, respectively (Figure 6C). In contrast, the marked reduction of uNK cells, as measured by as measured by tissue uNK cell proportion and normalized mean gland CD56 immunoreactivity, in RMM cases was not time-dependent (Figure 6C).

To determine whether these coordinated glandular and immune alterations collectively define a reproducible histological signature of RMM, a supervised random forest classifier incorporating glandular and cellular features was trained to distinguish RMM from control subjects and non-phenotyped miscarriage cases. During cross-validation, the model achieved an area under the receiver operating characteristic curve (AUROC) of 0.70 ± 0.01, effectively distinguishing 104 RMM patients from 3,238 other cases (Figure S9A). Consistent with the regression analyses, SHAP analysis highlighted glandular features (mean gland solidity, tissue lumen proportion) and uNK cell measurements (tissue uNK cell proportion and mean gland CD56 immunoreactivity) as the dominant predictors of RMM (Figure S9B).

### Endometrial glandular metabolic impairment associated with recurrent missed miscarriage

To further investigate Endometronome’s prediction that RMM is associated with a distinct pathological endometrial state, we performed single-cell RNA-sequencing on timed endometrial samples (LH+5 to +9 days) collected from control subjects (n = 7), women with a history of one or more bleeding-associated pregnancy losses (n = 10), and RMM patients (n = 4) (Table S1). Dimensionality reduction identified eight major cell clusters, comprising 8,509 uNK cells and 26,744 epithelial cells (Figure S10A). All samples contributed to glandular, ciliated, and luminal epithelial subpopulations, and, consistent with previous studies, four uNK subsets were identified: uNK1, uNK2, uNK3, and proliferative uNK cells (Figure S10A). Based on the morphological signature of RMM, we first explored epithelial and uNK cell crosstalk across patient groups using MultiNicheNet^47^, a framework for differential cell-cell communication analysis from multi-sample, multi-condition, single-cell transcriptomics data. MultiNicheNet analysis identified differentially active epithelial-uNK ligand-receptor interactions across control, miscarriage, and RMM conditions. Circos plots of the top 50 ranked ligand–receptor interactions revealed an overrepresentation of bidirectional epithelial-uNK cell receptor-ligand interactions in RMM compared with other miscarriage patients. Except for two, none of the predicted interactions associated with miscarriage featured prominently in control subjects (Figure 7A). To investigate the downstream consequences of these altered interactions, top-ranked ligand–receptor pairs were linked to downstream target genes to construct a regulatory signalling network. Filtering for ligand–target relationships with consistent expression correlation across samples yielded a high-confidence network specific to RMM (Figure 7B), predicting epithelial dependency on receptor tyrosine kinase (RTK) signaling downstream of EGFR and IGFR activation.

**Figure 7.**
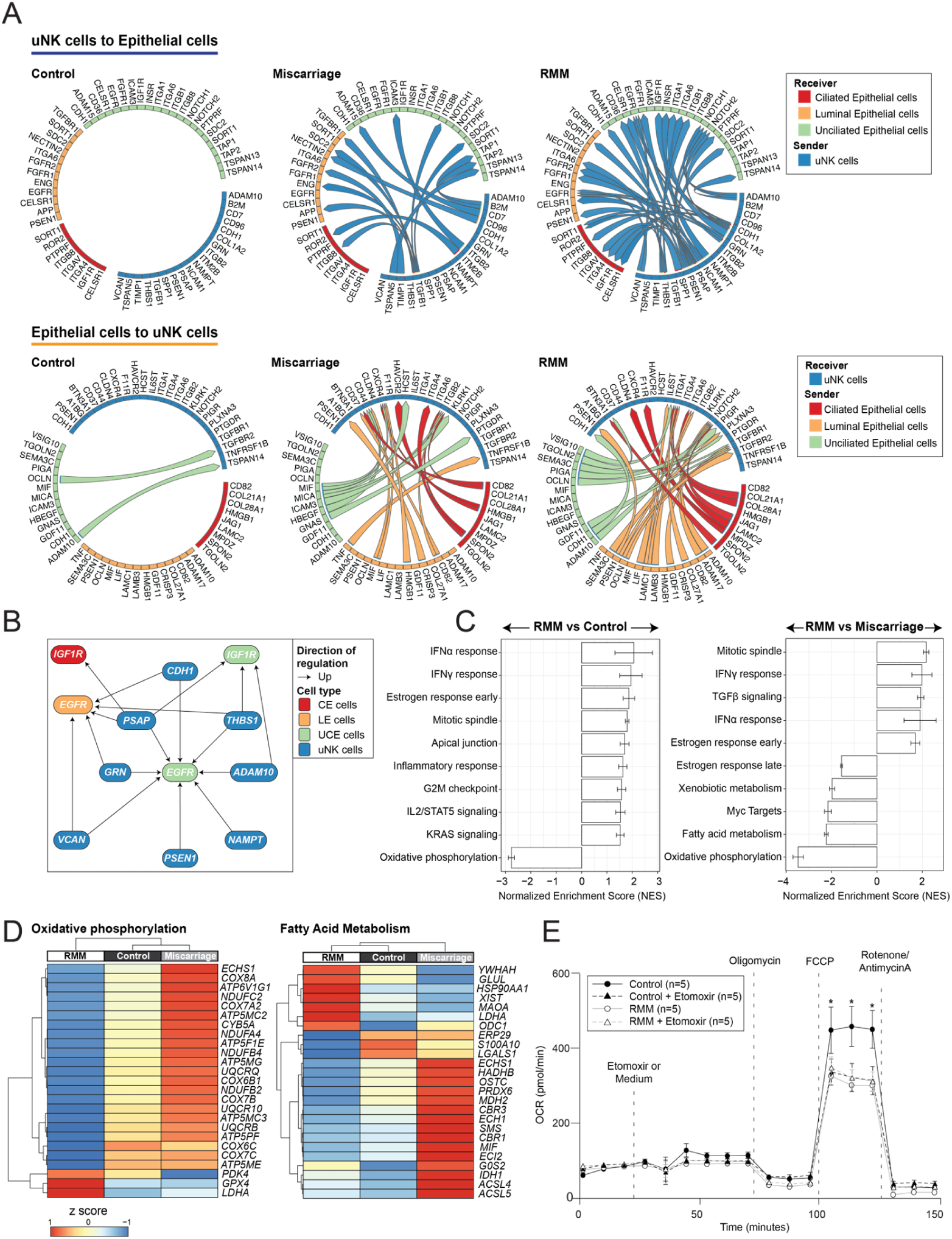
Recurrent missed miscarriage is associated with altered epithelial–uNK cell communication and impaired glandular metabolism. (A) Circos plots of the top-ranked epithelial–uNK ligand–receptor interactions identified by MultiNicheNet analysis. (B) MultiNicheNet filtered intercellular regulatory network showing prioritised ligand–target interactions between uNK and epithelial cell populations following ligand–target expression correlation filtering. (C) GSEA of Hallmark pathways in endometrial epithelial cells comparing RMM versus Control and RMM versus RPL epithelial cells. Bars show normalised enrichment scores (NES) for the top pathways, with error bars representing the standard error of NES across leave-one-RMM-donor-out analyses to indicate robustness. (D) Heatmaps showing row-scaled expression of the top 25 most variable genes in the Oxidative Phosphorylation, Fatty Acid Metabolism GSEA Hallmarks across RMM, Control, and RPL epithelial cells. (E) Mito Stress test OCR profile of intact organoids derived from 5 control (cyan) and 5 RMM (pink) endometrial biopsies. Points represent mean ± S.E.M. For each well, data were normalised to the third baseline reading.

Given that uNK cells are replenished from circulating progenitor cells in cycling endometrium^15^, the altered cell-cell interactions inferred by MultiNicheNet analysis likely reflect intrinsic glandular differences between patient groups. Hallmark gene set enrichment analysis (GSEA) was subsequently used to interrogate the epithelial cell fractions in the different patient groups. RMM epithelium was enriched for inflammatory and immune-associated pathways, including IFN-α and IFN-γ signaling, as well as pathways governing proliferation and structural organization (Figure 7C). In contrast, control epithelium was enriched for oxidative phosphorylation, consistent with a heightened metabolically oxidative state (Figure 7C and 7D). Direct comparison with the epithelial cells from non-RMM miscarriage patients also revealed enrichment of metabolic pathways in glandular epithelium in the non-RMM miscarriage group, including oxidative phosphorylation and fatty-acid metabolism (Figure 7C and 7D). Collectively, these findings suggest that RMM epithelium adopts an inflammatory state, accompanied by mitochondrial respiration suppression and a shift toward glycolysis, consistent with a Warburg-like metabolic program and in keeping with the inferred RTK dependency (Figure 7B)^48,49^.

To functionally validate these inferred metabolic alterations, we established endometrial gland-like organoids from RMM (n = 5) and control (n = 5) subjects (Table S4). Metabolic profiling via oxygen consumption rate (OCR) was performed following four days of culture, matrix recovery, and plating onto gel-coated microplates (Figure 7E). While basal and ATP-linked respiration were comparable between groups, maximal respiratory capacity was significantly reduced in RMM organoids, resulting in decreased spare respiratory capacity (*p* < 0.05, multiple Mann-Whitney test) (Figure 7E and Figure S10C). Inhibition of fatty-acid β-oxidation with etomoxir reduced spare respiratory capacity in control organoids to levels observed in RMM, but had no effect in RMM organoids. These findings indicate that reduced metabolic flexibility, reflecting impaired β-oxidation and increased dependency on glycolytic metabolism, in endometrial glands prior to conception may compromise histotropic nutrition in pregnancy, thereby increasing the risk of missed miscarriages.

## Discussion

Eutherian (placental) mammals utilize a wide range of strategies to establish a stable maternal-fetal interface during pregnancy^50,51^. This evolutionary diversity is attributed to coopetition, a game theory concept describing the balance of cooperative and competitive interactions between the genetically distinct embryo and the maternal endometrium^52^. Human embryos are characterized by genomic instability. Meiotic errors in oocytes account for the age-dependent increase in fetal aneuploidy, while postfertilization mitotic errors are age-independent and lead to embryonic mosaicism^53,54^. In this context, maternal control of the decidual reaction, which imposes a bistable state on the endometrium, serves as a critical countermeasure, as implantation of low-fitness embryos disrupt the decidual reaction and trigger menstruation-like tissue breakdown^3,11,55^. As a result of these evolved traits, human fecundity is relatively low, conception outcomes are inherently uncertain, and attrition rates in early gestation are high, with only a subset presenting as clinical miscarriages^3,56^. Nevertheless, when considered across over hundreds of potential conception cycles, this reproductive strategy is also adaptable and resilient, as evidenced by high cumulative live birth rates even after multiple pregnancy losses, particularly in younger women.

The shift in modern societal dynamics, characterized by a trend toward delayed childbearing, has significant reproductive implications, most prominently lower fertility and higher miscarriage rates, underscoring the need for novel diagnostics and therapeutics. Currently, miscarriage prevention strategies focus mainly on medical risk factors. However, the impacts of comorbidities and lifestyle factors on miscarriage rates are modest, and evidence-based treatments are, in most instances, not available^27,57,58^. Animal experiments have shown that endometrial perturbations at implantation induce adverse ripple effects, resulting in defective feto-placental development and poor pregnancy outcomes^59^. The accessibility of the human endometrium and the cyclicity of the decidual reaction facilitate the development of pre-pregnancy endometrial diagnostics and therapeutics to prevent pregnancy-related disorders. However, current methodologies, such as conventional histological dating and molecular assays, either fail to comprehensively assess the inflammatory decidual reaction or are too costly for clinical use, underscoring the need for scalable, comprehensive alternatives.

In this study, we present Endometronome, a heterogeneous graph neural network framework that robustly predicts endometrial dating, compartmental timing of the decidual reaction, biomarker ratios, and gene expression directly from digitised histology images. By integrating multi-scale features from image patches, glands, and individual nuclei, our model generates a comprehensive representation of tissue architecture and achieves consistently high predictive performance, surpassing self-reported timing in both cross-validation and independent test cohorts. Our results confirm that histological images contain substantial clinically relevant hidden information, as demonstrated by Endometronome’s capacity to quantify the impact of prior pregnancy losses on the decidual reaction, a proxy for recurrence risk^28,60^, through consolidative analysis of gene and biomarker predictions, glandular and nuclear parameters, community state, and whole-tissue features. Importantly, Endometronome revealed persistent perturbations in the decidual reaction associated with recurrent pregnancy loss that remain detectable across independent menstrual cycles. The framework also identified a distinct histomorphological signature of recurrent missed miscarriage, leading to the discovery of associated glandular metabolic impairment.

A major strength of Endometronome is its ability to combine predictive performance with spatial interpretability. Unlike conventional bulk molecular assays that generate a single measurement per biopsy, the framework produces localized spatial predictions across entire WSIs, enabling visualization of heterogeneous tissue states and compartment-specific remodeling dynamics. Feature attribution analyses further identified the glandular and nuclear structures most strongly associated with molecular and clinical predictions. This level of interpretability is particularly important for clinical deployment, where transparent reasoning and spatially grounded outputs are increasingly recognized as essential for trust, regulatory acceptance, and integration into diagnostic workflows^61–63^. More broadly, our findings demonstrate how AI models trained on routine histology can serve not only as predictive tools but also as discovery platforms capable of uncovering previously unrecognized biological states and mechanisms.

This study has several limitations. First, the cohorts were derived from an experimental reproductive medicine research clinic, potentially introducing biases and limiting generalizability to broader patient populations and clinical settings. To address this, future studies should validate the model prospectively using diverse patient populations, varying staining protocols, and subsequent pregnancy outcomes. Second, while our interpretability analyses reveal critical insights, the precise biological mechanisms underpinning features linked to miscarriage risk require further studies. Addressing these aspects will enhance clinical utility and biological understanding of our approach.

In summary, Endometronome establishes a scalable framework for decoding the spatial and molecular dynamics of the human decidual reaction directly from routine histology. By linking tissue architecture with molecular timing and reproductive history, the framework provides new insight into the biology of miscarriages while offering a potential foundation for pre-pregnancy diagnostics in reproductive medicine. More broadly, this work highlights the capacity of multiscale AI approaches to transform histological images into quantitative maps of tissue function, enabling both mechanistic discovery and clinically actionable assessment from standard pathology workflows.

## RESOURCE AVAILABILITY

### Data and code availability

All data needed to evaluate the conclusions in the paper are present in the paper and/or the Supplementary Materials. All data included in the study are provided in data files S1 and S2.

## ACKNOWLEDGMENTS

We thank the patients attending the Implantation Research Clinic, a dedicated experimental reproductive medicine clinic at University Hospitals Coventry and Warwickshire (UHCW) National Health Service (NHS) Trust, Coventry, UK. We also thank the staff of the Biomedical Research Unit in Reproductive Health, Warwick Innovation, Arden Tissue Bank, Institute for Precision Diagnostics and Translational Medicine, and Research and Development Department at UHCW NHS Trust. We are indebted to S. Quenby, A. Hawkes, J. Odendaal, L. Lacey, L. J. Ewington, S. Tewari, I. Pavlidis, M. Alkhoury, and E. Scott for assisting with sample collection and processing. We thank K. Fishwick, N. M. Adwani, Sean James, and J. Thornton for technical support and S. Ott for bionformatics advice. We thank Ethar Alzaid for valuable assistance in the design and preparation of figures. We are also grateful to the Simon Tandi and Skiros Habib for their support in setting up and maintaining the webserver.

## AUTHOR CONTRIBUTIONS

Conceptualization: J.J.B. and F.M. Funding acquisition: F.M. and J.J.B. Investigation: G.W., T.M.R., A.E., P.B., W.T.M.F., A.Y., K.M., H.Y., C.-S.K., M.C., E.S.L., and J.M. Methodology: J.D.A., F.M., J.J.B., M.E., G.W. and T.M.R. Visualization: J.M., M.T., G.W., and T.M.R. Supervision: F.M. and J.J.B. Writing-original draft: G.W., T.M.R., and J.J.B. Writing-review and editing: J.M., A.E., P.R.B., and F.M.

## DECLARATION OF GENERATIVE AI AND AI-ASSISTED TECHNOLOGIES

During the preparation of this work, the author(s) used perplexity to improve spelling and grammar. After using this tool or service, the author(s) reviewed and edited the content as needed and take(s) full responsibility for the content of the publication.

## METHODS

### Study Participants and Ethical Approval

This study was conducted at the Implantation Research Clinic, a dedicated experimental reproductive medicine clinic at University Hospitals Coventry and Warwickshire (UHCW) National Health Service (NHS) Trust, Coventry, UK. Patients attending the clinic for endometrial assessment presented with a history of one or more miscarriages and/or fertility issues, including unexplained, male factor, and tubal infertility. We defined miscarriage as the spontaneous loss of pregnancy up to 24 weeks of gestation, excluding pregnancies of unknown location, ectopic or molar pregnancies, and terminations for any reason. We adopted a broad definition of miscarriage encompassing all spontaneous pregnancy losses without a requirement for ultrasound confirmation in line with the recommendations from an international consensus development study on core outcome sets for miscarriage studies^64^. Surplus endometrial samples obtained for diagnostic purposes were processed for research. The use of surplus samples for research was approved by the Arden Tissue Bank at UHCW NHS Trust (NHS Research Ethics Committee approval: 23/WA/0182). All samples were obtained following written informed consent and in accordance with the Declaration of Helsinki (2000) guidelines.

### Endometrial Sample Collection

Anonymized endometrial samples were obtained during the luteal phase of ovulatory menstrual cycles and timed relative to the pre-ovulatory LH surge, as determined by commercially available ovulation test kits. Overt uterine pathology was excluded by vaginal ultrasonography scan prior to the biopsy. Endometrial biopsies were obtained using a Wallach Endocell™ or CerviX™ endometrial sampler. Paired endometrial biopsies were obtained in 943 study participants. Two or more regular menstrual cycles were required for repeat endometrial sampling following a pregnancy loss. A minimal set of anonymized demographic data was collected for each endometrial sample (Table S1, Table S3, and Table S4).

### CD56 Immunohistochemistry

Formalin-fixed endometrial biopsies were processed through graded alcohol and xylene using a Shandon Excelsior ES Tissue Processor (ThermoFisher Scientific, Paisley, UK), before paraffin-embedding in Surgipath® tissue infiltration medium (Leica Biosystems, Wetzlar, Germany) using a Tissue-Tek TEC embedder (Sakura Finetek Europe, Netherlands). Formalin-fixed, paraffin-embedded (FFPE) endometrial biopsies were sectioned at 3 µm using a microtome and mounted on Surgipath Xtra slides (Leica Biosystems) by overnight incubation at 60 °C. CD56 immunostaining and DAB detection were fully automated on a Bondmax autostainer (Leica Biosystems), using sodium citrate buffer for antigen retrieval (10 mM sodium citrate, 0.05% Tween-20, pH 6). Sections were stained with anti-CD56 (NCAM-1) antibody (Novocastra NCL-L-CD56-504, 1:200; Leica Biosystems) and counterstained with haematoxylin. Human tonsil tissue on each slide served as a positive control. Negative controls were included by omitting the primary antibody on human tonsil and high-CD56 control biopsies, and staining runs were repeated if control results were not as expected. Slides were dehydrated, cleared, and cover-slipped using a TissueTek Prisma Automated Slide Stainer (Model 6134; Sakura Finetek Europe) with DPX mountant. Bright-field images were acquired using a Mirax Midi slide scanner (Zeiss, Oberkochen, Germany) with a 20× objective and at a spatial resolution of 0.50µm per pixel, and saved as mrxs files to visualise and for analysis.

### RT-qPCR Analysis

Total RNA was extracted from RNAlater-protected endometrial biopsies using STAT-60 (AMS Biotechnology, Oxford, UK), according to the manufacturer’s instructions. Reverse transcription was performed from 1 µg RNA using the Quantitect Reverse Transcription Kit (QIAGEN, Manchester, UK) and cDNA was diluted to 10 ng/µl equivalent before use in qPCR. Amplification was performed on a Quant5 Real-Time PCR system (Applied Biosystems, Paisley, UK) in 10 µl reactions using 2× QuantiFast SYBR Green PCR Master Mix containing ROX dye (QIAGEN), with 300 nM each of forward and reverse primers. *L19* was used as a reference gene. Primer sequences have been described previously^41^.

### Data Acquisition and Preprocessing

In this study, the majority of our WSIs contained control tissue from the small bowel alongside the target endometrial tissue. To ensure accurate analysis, we employed a user-defined area selection method to precisely identify the boundaries of the endometrial tissue. This approach simultaneously allowed us to eliminate artefacts such as staining errors and pen marks. Within the defined area of interest, we applied a threshold of 250 on the average pixel intensity, effectively segregating the tissue regions from the background. For patch graphs, we tiled the tissue region into 224×224 pixel patches, using an image resolution of 0.50 mpp. To ensure the quality and relevance of our data, we implemented a filtering process for the generated patches. We excluded patches with less than 5% informative tissue area, where informative area was determined by pixels with a mean intensity greater than 220. Any remaining patches were included for analysis.

### Detection of Histological Components

We processed the tissue area of each WSI using the ASTER model, a convolutional neural network (CNN)-based segmentation model specifically designed for endometrial tissue analysis^65^. ASTER employs a shared ResNet34 encoder, to efficiently extract features from input images, and distinct U-Net style decoders for each annotation class. This enables an efficient simultaneous segmentation of multiple endometrial structures. The model was trained on a dataset of 570 annotated image patches, encompassing 35,135 objects across six histological classes, and has demonstrated high segmentation accuracy and generalizability. For this study, we specifically utilized the segmented outputs corresponding to glands and nuclei as the basis for subsequent feature extraction and graph construction. The application of ASTER to our dataset enabled the identification and localization of 1,186,835,994 nuclei and 2,644,514 gland components, along with 9,296,962 image patches providing a comprehensive and multi-resolution foundation for graph-based modelling of endometrial architecture. Figure 1 shows example areas of segmented glands and nuclei used in our analysis.

### Feature Extraction

Our feature extraction strategy is tailored to the unique characteristics of each node type within the heterogeneous graph, integrating both deep learning-derived representations and clinically interpretable attributes. This dual approach ensures that the resulting node features capture complex visual patterns while retaining interpretability.

### Patch Representation

For patch nodes, each representing a 224 × 224 pixel region of the tissue, we employed a deep feature extraction pipeline using ShuffleNet V2^66^, a lightweight convolutional neural network pretrained on ImageNet^67^. Features were extracted from the penultimate fully connected layer, yielding a 1024-dimensional embedding *h_i_* ∈ ℝ^U024^ for each patch node. This high-dimensional representation encodes visual information, capturing morphological and textural patterns within each tissue region.

### Nuclei Representation

For nuclei nodes, we derived a set of 15 interpretable features encompassing colour and morphology characteristics. Colour features included the mean, maximum, median, and standard deviation of the hematoxilyn channel and CD56 channels within the bounding area, quantifying staining intensity and variability. Morphological descriptors comprised area, solidity, eccentricity, circumference, equivalent diameter, major and minor axis lengths, and convex area of the nuclei boundary contour. Collectively characterizing nuclei size and shape. These features provide an informative summary of nuclear phenotypes within the tissue. Figure S4 shows a visual representation of the features used with descriptions included in Table S5.

### Gland Representation

Gland nodes were characterized by an expanded feature set, incorporating both the colour and morphological features described for nuclei, as well as additional attributes capturing gland-cell relationships and epithelial organization. Specifically, we computed the number of constituent nuclei, nuclei density (total nuclei number divided by gland area), and statistics (mean, median, maximum, standard deviation) of distances from epithelial nuclei centroids to their nearest gland boundary. To further quantify epithelial pseudo stratification, we calculated the number of neighbouring epithelial nuclei within a 50-pixel radius and the distances to the eight nearest neighbours. This comprehensive feature set resulted in a 29-dimensional vector for each gland node, capturing both intrinsic gland properties and their cellular microenvironment (Figure S4 and Table S6). To ensure comparability across node types and mitigate scale differences, all nuclei and gland features were normalized using z-scoring. This systematic feature engineering approach supports the integration of both deep features and domain-specific information in the heterogeneous GNN framework.

### Graph Representation

To accurately represent the multi-scale architecture of endometrial tissue, we constructed a heterogeneous graph *G* = (*V*, *E*, *T_V_*, *T_E_*) from each segmented WSI. In this graph, each node *v_i_* ∈ *V* corresponds to an entity, a nuclei, gland, or image patch, and is assigned a node type *n* ∈ *T_V_*, where *T_V_* = {nuclei, gland, patch}. Each node *v_i_* ∈ *V* is therefore characterized by a tuple (*n_i_,* **g***_i_,* **h***_i_*), where **g***_i_* denotes the spatial coordinates of the centroid of the object, and **h***_i_* is its feature vector as described in Section 4.5. The edge set *E* is constructed separately for each biologically relevant edge type Specifically, we defined five edge types *T_E_* = {patch-patch, nuclei-nuclei, gland-gland, gland-patch, and nuclei-patch}, thereby capturing both direct spatial interactions among similar tissue components and hierarchical associations across biological scales. Hyperparameters were selected empirically to balance local tissue context with computational tractability.

### Homogeneous Subgraph Construction

Edges *e* ∈ *E*, where *e* = (*v_i_*, *v_j_*, *r*), are instantiated between nodes v*_i_* and v*_j_* based on spatial proximity and biological context, with each edge annotated by type *r* ∈ *T_E_*. For patch-patch and gland-gland edges, we employ Delaunay triangulation among nodes within a 3,000-pixel Euclidean distance, resulting in planar subgraphs. For nuclei-nuclei edges, each nuclei node is connected to its 9 nearest neighbours within a 3,000-pixel threshold, due to the computational complexity of including more neighbours, forming a non-planar subgraph.

### Heterogeneous Graph Representation

To integrate these subgraphs into a unified heterogeneous structure, we introduce edges between node types. Nuclei-patch edges are established by linking each nuclei node to the patch node in which the centroid of the nuclei resides. Similarly, gland-patch edges are formed by connecting each gland node to any patch node with which it shares overlapping pixels. By explicitly modelling these cross-level edges, the unified graph structure integrates tissue architecture across multiple spatial scales, from fine-grained nuclear interactions to larger glandular and tissue-level patterns. This multi-scale representation captures the complex spatial relationships inherent in biological tissues, thereby supporting more comprehensive and biologically in-formed analyses. Figure 1 illustrates the overall workflow for generating the heterogeneous graphs.

### Heterogeneous Graph Neural Network Architecture

The heterogeneous graph representations constructed from each WSI served as input to a heterogeneous graph neural network (HGNN) designed to predict the 16 WSI-level target variables, EndoTime, LH+, two gene ratios, and 12 gene expression values. Our HGNN architecture leverages different graph convolutional operations for distinct edge types, enabling specific feature propagation and effective modelling of the hierarchical organization present in endometrial tissue. Figure S2A provides a visual summary of the model architecture.

### Message Passing and Edge-Type-Specific Operations

Within this framework, node features are iteratively updated through a sequence of two message-passing layers, each tailored to the biological context and connectivity of the graph. For nodes representing glands and patches, we employed EdgeConv layers^68^, which are well-suited for capturing local geometric relationships. For the densely connected nuclei subgraphs, we utilized Graph Isomorphism Network (GIN) layers^69^, which provide strong discriminative power and computational efficiency for large-scale graphs. This approach allows the model to flexibly combine information from each node and its neighbours, facilitating the learning of complex patterns within the cellular microenvironment.

### Multi-Level Message Passing

To further capture higher-order interactions and multi-level relationships, we introduced additional message passing layers edges: GIN layers for nuclei–patch edges and EdgeConv layers for gland–patch edges. These layers enable information exchange between different biological scales, allowing the model to integrate fine-grained nuclear features with broader tissue context. At each stage of the backbone pipeline, these cross-level operations update the patch-level features for readout only, while the original graph remains unchanged for subsequent layers. This design ensures that the readouts reflect the evolving state of the graph. Although supervision was available only at the biopsy level, the graph architecture enabled inference of spatially localized tissue states through propagation of multiscale contextual information across the WSI.

### Patch-Level Readouts and WSI-Level Prediction

At each graph layer, patch-level predictions are obtained by applying a multi-layer perceptron to the feature representations of the patch nodes. These patch-level scores are then aggregated using mean pooling to yield a WSI-level prediction for that layer:

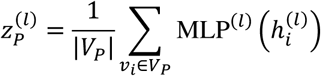

Where 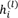 denotes the feature vector of patch node *v_i_* at layer *l*, V_P_ is the set of patch nodes, and MLP^(*l*)^ is the layer-specific multi-layer perceptron.

The WSI-level predictions from each layer are then summed to produce the final graph-level output:

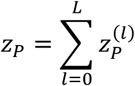

This iterative, multi-scale architecture, as visualized in Figure S2A, enables progressive integration of local nuclear and glandular contexts with global tissue patterns. Each refinement step enhances the model’s capacity to capture spatially nested relationships, ultimately supporting biologically informed WSI-level predictions.

### Training and Performance Evaluation

Model parameters, including those of the EdgeConv and GIN layers as well as the node-level MLPs, are optimized using backpropagation with a pairwise ranking loss. For a batch of size N and target variables K, the loss is defined as:

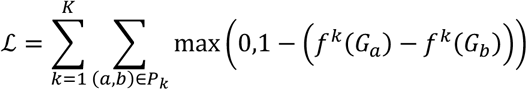

Where *f^k^*(*G*) is the prediction of variable *k* for graph *G* and 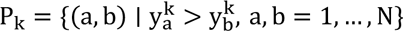 is the set of all sample pairs in the batch for which the true value of *k* is greater for *a* than for *b*. This ranking loss encourages the model to learn representations that preserve the correct ordering of samples with respect to each clinical variable. We employed a pairwise ranking objective rather than direct regression to prioritize preservation of biologically meaningful temporal and molecular ordering across samples while reducing sensitivity to inter-patient variability and scaling differences between targets.

We evaluated our HGNN model using a rigorous 5-fold cross-validation framework. In each fold, 80% of the data was allocated for training and 20% for testing. Within the training set, 10% was further set aside for validation and hyperparameter tuning. Model training was conducted for up to 300 epochs with a learning rate of 0.001 and weight decay of 0.0001. The Adam optimizer was employed^70^, using a batch size of 8. To prevent overfitting, early stopping was implemented, terminating training if validation performance did not improve for 20 consecutive epochs. For the final independent test set inference, we averaged the models generated from the 5 folds to produce final prediction scores for each instance.

As our ranking loss function yields prediction scores that are not directly comparable to ground truth values, we applied min-max scaling to normalize model outputs to the range of the ground truth for each variable. This normalization was particularly important for timing estimates, where raw prediction scores lack direct interpretability. Model performance was assessed using Spearman’s rank correlation coefficient (ρ) between predicted and observed values. We report the mean and standard deviation of ρ across the five folds, as well as the corresponding p-values to evaluate statistical significance. Predictions on the test set are bootstrapped to give confidence intervals and standard deviations.

### Graph Explainability

To interpret the decision-making process of our HGNN, we employed the GNNExplainer framework for heterogeneous GNNs^71^. This approach optimizes a feature or node mask that assigns an importance value, highlighting which features and nodes are most significantly influencing the model’s predictions. The feature mask values range between 0 and 1, with higher values indicating greater relevance. All explainability analyses were performed on representative samples from the test set to ensure that the explanations reflect the model’s generalization behavior.

### Timing Normalization

Miscarriage-associated analyses were performed on timing-normalized predictions to reduce confounding by cycle stage and isolate deviations from expected decidual progression. For each LH+ day, feature values were normalized using parameters estimated from day-specific fitted normal distributions. These fitted distributions were then used to convert each observation to its cumulative percentile for interpretability. This approach reduced confounding by cycle stage and enabled comparison of features relative to the expected distribution for each day of the cycle.

### Generalised Linear Model Analysis

Machine learning–derived predictors and interpretable histological features were assessed for association with number of previous miscarriages using Bayesian generalized linear models (GLMs) using a negative binomial regression model (GLM with a log link). Weakly informative priors were placed on model parameters. Intercepts were assigned a Normal distribution with mean zero and standard deviation of 10. In the negative binomial models, an additional dispersion parameter α was introduced with a Half-Normal prior distribution with standard distribution of 1. For the slopes of all predictors, we specified a Normal distribution with mean zero and standard deviation of 0.5, reflecting regularization at the scale of a 1% increase in the predictor. Posterior estimates were summarized using posterior medians and 90% credible intervals derived from the highest density intervals (HDIs). Evidence for an association between predictors and outcomes was considered present when the credible interval for the incidence rate ratio (IRR), calculated as the exponentiated regression coefficients (/3), excluded 1. This criterion reflects the probability the true IRR lies entirely above or below the null value, indicating a statistically meaningful effect.

### Differential Community Abundance Analysis

We analyzed all 2,644,514 gland components, each represented by a 29-dimensional feature vector. Z-score normalized features were embedded into two dimensions using Uniform Manifold Approximation and Projection (UMAP) to preserve local and global structure^72^, followed by k-means clustering to identify 10 distinct morphological communities. Each gland component was assigned to a single cluster, which was subsequently treated as a community membership identifier for downstream analyses. To ensure sufficient model degrees of freedom, community-level analyses were restricted to nine clusters, excluding one reference cluster.

To test whether community abundances varied with clinical outcomes (timing and number of previous miscarriages), we performed differential community abundance analysis using DCAFA^73^. For each community *k* and patient *i*, community counts c_ik_ were modelled using negative binomial regression with bag size offset:

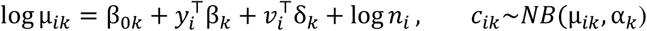

Where μ_ik_ is the expected count α*_k_* is a dispersion parameter, y_i_ represents clinical target variable, *v_i_* denotes covariates, and *n_i_* is the number of glands per patient. Exponentiated coefficients exp(β*_k_*) quantify fold-changes in community abundance per unit change in outcome. Significance was assessed using likelihood ratio tests with false discovery rate (FDR) correction across communities.

### Prediction of RMM

To assess the predictive value of interpretable features for RMM, we constructed a classification framework based on quantitative features derived from segmentations of WSIs. For each patient, a 13-dimensional feature vector was generated from day normalised WSI mean features, including solidity, eccentricity, area, CD56, and Hematoxylin staining of gland and nuclei. In addition, we included the tissue lumen proportion, the area of lumen divided by the area of the tissue mask, and uNK cell proportion, number of nuclei with CD56 staining over a threshold, as tissue level descriptors. Also included were clinical variables (age, BMI, and LH+ day). Classification was performed using a random forest algorithm implemented with balanced class weights, a minimum of four samples required to split an internal node, a minimum of 4 samples per leaf, and an ensemble of 200 decision trees. These were the optimal hyperparameters selected to mitigate overfitting. Model training was conducted to distinguish between patients with RMM and those without. Feature importance was assessed using SHAP analysis^74^, facilitating the identification of the most discriminative glandular attributes. Model training and evaluation employed stratified 5-fold cross-validation to maintain balanced class distributions. Predictive performance was assessed using the area under the receiver operating characteristic curve (AUROC) metric.

### Processing Endometrial Samples for scRNA-seq

Endometrial biopsies in additive-free DMEM/F12 media were processed within 15 min of collection. Samples were mechanically minced and enzymatically digested with DNase I (100 µg/mL), Collagenase Type Ia (500 µg/mL) and IV (10 µg/mL) for 90 min at 37°C, with shaking every 15 min. Undigested tissue was removed through a 40 μm cell strainer. To eliminate red blood cells, the cell suspension was incubated with red blood cell lysis buffer (Invitrogen, 00-4333-57) for 5 min at room temperature. The reaction was terminated by adding 30 mL PBS and centrifuged at 400 × g for 5 min. Cells were then processed according to Demonstrated Protocol CG00054 (Revision B, 10X Genomics) with some modifications. Briefly, cells were centrifuged at 300 × g for 5 min and resuspended in 0.04% BSA in PBS. This was repeated twice more before cells were resuspended in 500 μL PBS with 0.04% BSA and counted. Cells were adjusted to 700 cells/μL.

### scRNA-seq Data Processing and Annotation

Single-cell RNA-seq data comprising 143,597 cells from 21 luteal phase endometrial biopsies were analysed using the Seurat framework in R. Cells were grouped by patient phenotype and day post–luteinizing hormone (LH) surge for downstream analyses. Dimensionality reduction was performed using principal component analysis followed by Uniform Manifold Approximation and Projection (UMAP) to visualise cellular heterogeneity. Cells were broadly classified into stromal, epithelial, immune, and endothelial compartments based on canonical marker gene expression. These clusters were subsetted for further analysis and reclustered, revealing distinct subpopulations defined based on established marker genes and transcriptional profiles.

### Ligand-Receptor Analysis

Cell–cell communication between uNK cells and epithelial populations was analysed using the MultiNicheNet framework in R^47^. Analyses were performed across clinical groups while accounting for sample-level structure. In brief, ligand-receptor interactions were prioritised and top-ranked interactions were identified across conditions and visualised using circos chord diagrams to summarise dominant sender-receiver communication patterns. To further characterise downstream signalling, ligand–target links were extracted and used to construct intercellular regulatory networks, representing ligands, receptors, and inferred target genes as interconnected nodes. Networks were generated both with filtering for ligand–target expression correlation across samples; correlation-filtered networks (Pearson correlation threshold applied) were used for primary interpretation as they are more likely to reflect active signalling relationships. Additional visualisations, including bubble plots, were used to decompose prioritisation scores and facilitate interpretation of the underlying evidence supporting each interaction. Quality control assessments included evaluation of the number of cells per cell type per sample and inspection of statistical robustness of differential expression signals (adjusted *p*-values), ensuring that prioritised interactions were not driven by cell number imbalances or low-confidence transcriptional changes.

### Gene Set Enrichment Analysis

Differential gene expression analysis was performed on the epithelial subset using the Seurat function FindMarkers, comparing recurrent miscarriage (RMM) versus Control and RMM versus recurrent pregnancy loss (RPL) groups. Genes were ranked by average log2 fold change (avg_log2FC), excluding genes with missing values or duplicated gene symbols. Ranked gene lists were sorted in decreasing order and used as input for gene set enrichment analysis. Gene set enrichment analysis was conducted using the fgsea package (fgseaMultilevel function). Hallmark gene sets were obtained from the MSigDB Hallmark gene sets via the msigdbr package (category “H”, Homo sapiens). Enrichment was quantified using normalised enrichment scores (NES), and statistical significance was assessed using adjusted *p*-values (Benjamini–Hochberg correction). For visualization, the top-ranked pathways were selected based on adjusted p-value or defined a priori and plotted according to NES. To assess robustness to donor variability, a leave-one-donor-out approach was implemented for RMM samples. In each iteration, one donor was excluded, and differential expression and GSEA were repeated as described above. Pathway-level NES values across iterations were aggregated, and variability was quantified as the standard error of the NES, calculated from non-missing values across runs. These values were incorporated as error bars in GSEA visualizations derived from the full dataset.

### Pathway-Level Gene Expression Analysis and Heatmaps

Gene expression within selected Hallmark pathways was further examined at the gene level. Pathway-specific gene sets were extracted from MSigDB using msigdbr. Average gene expression across experimental groups (RMM, Control, and RPL) was calculated using the AverageExpression function in Seurat. Genes with zero variance across groups were excluded. For each pathway, the top variable genes were identified based on standard deviation across group-averaged expression values, and the top 25 genes were selected for visualisation. Heatmaps were generated using the pheatmap package. Expression values were scaled by row (z-score normalisation) to emphasize relative differences across conditions. Both genes and conditions were hierarchically clustered using default parameters. Column annotations were included to indicate group identity.

### Endometrial Gland Organoid Cultures

Fresh tissue samples were washed in a growth medium and finely minced using a scalpel. Minced tissue was digested for 1 hour at 37 °C with collagenase type IA (0.5 mg/ml) (Sigma-Aldrich) and deoxyribonuclease type 1 (0.1 mg/ml) (Lorne Laboratories, Reading, UK) in phenol red-free DMEM/F12 medium. The digested tissue was filtered through a 40 µm cell strainer and rinsed with 5 mL of 10% growth medium. Endometrial stromal cells and blood cells were collected through the flow-through, whereas glandular clumps were retained. For the isolation of glands, the strainer was inverted over a 50 ml Falcon tube and backwashed. Both cell types were centrifuged at 280 g for 5 minutes, and cell pellets were cryopreserved in 10% DMSO in dextran-coated charcoal-treated fetal bovine serum.

Organoids were generated from frozen isolated gland fragments of 5 control subjects and 5 RMM patients as described previously^75^. Samples were centrifuged and digested in 500 µl TrypLE Express Enzyme (1×) for 10 minutes in a shaking water bath. Enzyme was diluted with phenol red-free DMEM/F12, and the cell suspension was passed through a 40 µM strainer and centrifuged. The pellet was resuspended, and the cells were counted. Pellets were resuspended in ice-cold Matrigel at a concentration of 20 cells per µl. 5 µl droplets were plated in a 96-well plate and incubated at 37 °C for 15 minutes to cure the gel. A minimum of three wells were plated per each sample. Afterwards, 100 µl of expansion medium, as described previously, supplemented with 10 µM Y-27632 (Abcam) was added to each well for the first 3 days. Medium was replaced every 2-3 days.

### Seahorse XF Mito Stress Test Assay

All products for this assay were from Agilent (Cheadle, UK) unless otherwise stated. The Seahorse sensor cartridge was hydrated at 37 °C in the non-CO_2_ incubator overnight, following the manufacturer’s instructions. Day 4 gland organoids at passage 2 from control and RMM endometrial biopsies were removed from the gel using ice-cold PBS at 4 °C. During this time, 50 µl of gel diluted 1:10 in ice-cold Seahorse XF Assay Medium (Seahorse XF DMEM medium supplemented with 1 mM pyruvate, 2 mM glutamine, and 10 mM glucose, pH 7.4) was added to each well of the Seahorse XF Cell Culture Microplates excluding four blank wells and incubated for 1 hour at 37 °C. After 1 hour, the organoids were centrifuged at 300 g for 5 minutes, resuspended in fresh PBS, and centrifuged again. Samples were resuspended in Seahorse XF Assay Medium, counted, and resuspended at a density of 3 organoids per µl. The excess solution from the Seahorse XF Cell Microplate was aspirated and 50 µl of the sample suspension was evenly distributed in each well. The plate was incubated for an hour at 37°C to allow the organoids to settle. Each well was topped up with 400 µl Seahorse XF Assay Medium and the plate was placed in a 37 °C non-CO2 incubator for 45 minutes to equilibrate. Seahorse Assay medium, 1 µM oligomycin, 2 µM FCCP, 0.5 µM antimycin A, and 4 µM etomoxir were used following the manufacturer’s instructions. All samples were run with five replicates in Seahorse XF24 Analyser. The data were analysed in Wave, Microsoft Excel and GraphPad Prism, and normalised to the 3rd baseline measurement. Mann-Whitney test was used to test for comparison of two unpaired, non-parametric datasets.

## SUPPLEMENTAL INFORMATION

### TABLES

**Table S1.**
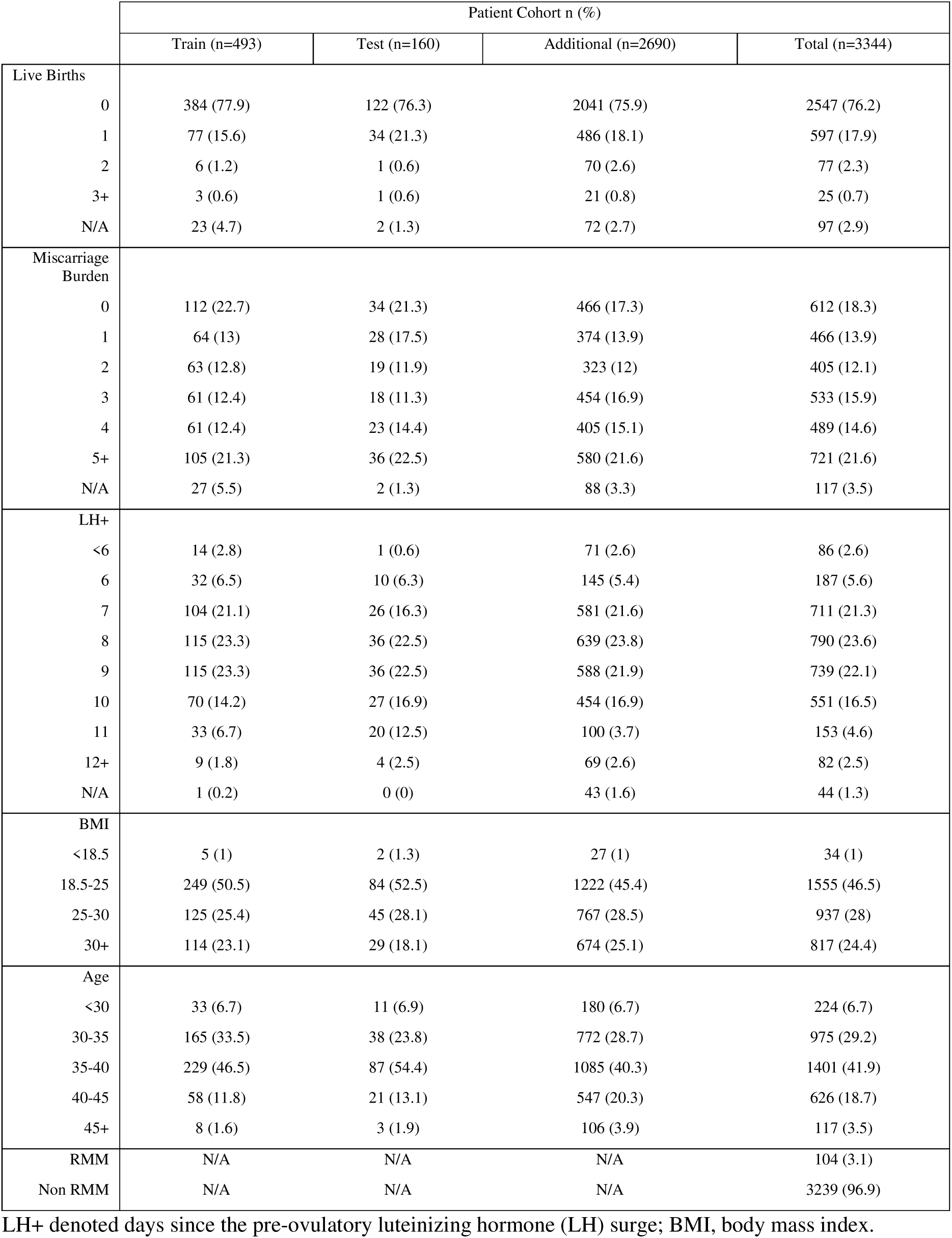
Demographic and clinical information for the patient cohort.

**Table S2.** Comparison of model performance using Spearman’s correlation coefficients across predicted variables.

|  | Patch CV | Nuclei CV | Gland CV | EndoMetronome CV | Patch Test | Nuclei Test | Gland Test | Endometronome Test |
| --- | --- | --- | --- | --- | --- | --- | --- | --- |
| *GPX3/SLC15A2* ratio | 0.821±0.024 | 0.714±0.055 | 0.817±0.023 | 0.819±0.036 | 0.721±0.051<br>(0.609-0.807) | 0.601±0.056<br>(0.486-0.702) | 0.727±0.055<br>(0.610-0.823) | **0.746±0.047**<br>**(0.646-0.827)** |
| EndoTime | 0.778±0.038 | 0.660±0.052 | 0.809±0.035 | 0.783±0.039 | 0.724±0.047<br>(0.629-0.807) | 0.587±0.055<br>(0.463-0.680) | 0.722±0.051<br>(0.613-0.811) | **0.737±0.047**<br>**(0.631-0.815)** |
| *DPP4* Expression | 0.768±0.045 | 0.654±0.045 | 0.784±0.044 | 0.770±0.055 | 0.722±0.051<br>(0.608-0.807) | 0.603±0.053<br>(0.491-0.688) | **0.744±0.049**<br>**(0.637-0.827)** | 0.741±0.048<br>(0.637-0.826) |
| *GPX3* Expression | 0.758±0.044 | 0.633±0.061 | 0.767±0.039 | 0.762±0.046 | 0.694±0.049<br>---(0.593-0.781) | 0.593±0.052<br>(0.488-0.689) | **0.743±0.047**<br>**(0.651-0.826)** | 0.735±0.044<br>(0.647-0.815) |
| *SLC15A2* Expression | 0.760±0.014 | 0.687±0.035 | 0.744±0.051 | 0.753±0.056 | 0.653±0.055<br>(0.531-0.746) | 0.538±0.067<br>(0.393-0.659) | 0.661±0.057<br>(0.546-0.760) | **0.666±0.056**<br>**(0.544-0.766)** |
| *CXCL14* Expression | 0.605±0.079 | 0.507±0.076 | 0.641±0.041 | 0.627±0.067 | 0.630±0.053<br>(0.524-0.719) | 0.415±0.070<br>(0.271-0.538) | 0.664±0.051<br>(0.556-0.756) | **0.673±0.051**<br>**(0.572-0.762)** |
| *ITGAD* Expression | 0.590±0.094 | 0.586±0.077 | 0.601±0.038 | 0.610±0.064 | 0.610±0.057<br>(0.492-0.717) | 0.581±0.060<br>(0.455-0.687) | 0.581±0.059<br>(0.458-0.691) | **0.631±0.055**<br>**(0.514-0.724)** |
| *IL2RB* Expression | 0.643±0.068 | 0.622±0.069 | 0.612±0.038 | 0.659±0.047 | 0.548±0.065<br>(0.410-0.665) | 0.521±0.065<br>(0.390-0.637) | 0.546±0.060<br>(0.421-0.653) | **0.606±0.060**<br>**(0.477-0.709)** |
| LH+ | 0.591±0.039 | 0.573±0.040 | 0.627±0.065 | 0.585±0.027 | **0.632±0.057**<br>**(0.512-0.729)** | 0.562±0.063<br>(0.436-0.675) | 0.624±0.062<br>(0.496-0.737) | 0.591±0.067<br>(0.444-0.707) |
| *DIO2* Expression | 0.626±0.043 | 0.500±0.044 | 0.618±0.061 | 0.600±0.060 | 0.474±0.066<br>(0.332-0.596) | 0.393±0.072<br>(0.251-0.528) | 0.517±0.062<br>(0.384-0.633) | **0.535±0.064**<br>**(0.407-0.648)** |
| *PLA2G2A/DIO2* Ratio | 0.462±0.063 | 0.504±0.034 | 0.606±0.089 | 0.605±0.057 | 0.487±0.066<br>(0.353-0.607) | 0.427±0.071<br>(0.283-0.558) | 0.483±0.068<br>(0.336-0.602) | **0.547±0.061**<br>**(0.421-0.653)** |
| *TIMP3* Expression | 0.501±0.066 | 0.448±0.087 | 0.488±0.079 | 0.489±0.095 | 0.419±0.067<br>(0.287-0.547) | 0.431±0.069<br>(0.293-0.563) | 0.355±0.072<br>(0.209-0.490) | **0.479±0.064**<br>**(0.349-0.598)** |
| *PLA2G2A* Expression | 0.609±0.061 | 0.386±0.043 | 0.479±0.117 | 0.455±0.092 | 0.397±0.069<br>(0.263-0.525) | 0.356±0.073<br>(0.206-0.487) | 0.383±0.069<br>(0.242-0.508) | **0.453±0.065**<br>**(0.318-0.574)** |
| *IL15* Expression | 0.315±0.070 | 0.261±0.118† | 0.276±0.053 | 0.317±0.059 | 0.345±0.074<br>(0.204-0.482) | 0.365±0.072<br>(0.218-0.496) | 0.337±0.074<br>(0.182-0.472) | **0.441±0.065**<br>**(0.300-0.561)** |
| *TAGAP* Expression | 0.238±0.131† | 0.232±0.062 | 0.201±0.095† | 0.238±0.108† | **0.320±0.070**<br>**(0.180-0.453)** | 0.255±0.077<br>(0.089-0.389) | 0.273±0.077<br>(0.106-0.412) | 0.308±0.075<br>(0.159-0.441) |
| *SCARA5* Expression | 0.230±0.062 | 0.047±0.137† | 0.234±0.107† | 0.203±0.064† | 0.158±0.077<br>(0.006-0.313) | 0.004±0.078<br>(-0.149-0.152)† | **0.203±0.081**<br>**(0.033-0.358)** | 0.120±0.079<br>(0.000-0.277)† |

Columns 2-5 report mean ± standard deviation from 5-fold cross-validation (CV), while columns 6-9 present independent test set performance with 95% bootstrap confidence intervals (CI) in parentheses. The proposed model is benchmarked against models utilizing only gland, nuclei, or patch subgraphs. All reported Spearman’s correlation $p$ -values are statistically significant ( $p < 0.05$ ) unless indicated by †.

**Table S3.** Demographic and clinical information for the patient cohort for Figure 7A-7D and Figure S10A and S10B.

| Group | n = | Age | LH+ (days) | Miscarriages | BMI |
| --- | --- | --- | --- | --- | --- |
| Total | 21 | 36.4 (33.5-39) | 7.2 (6-8) | 4.5 (1-6) | 25.5 (22-28.8) |
| Control | 7 | 38.17 (35.8–40.8) | 7.8 (6.8-8.3) | 0.5 (0-1) | 22.8 (20-26) |
| RMM | 4 | 35.5 (32.8-39) | 7.5 (5.8-8.8) | 5.8 (3.3-9) | 26.8 (22-29.8) |
| Miscarriage | 10 | 35.45 (32-41) | 6.9 (5-8) | 6.3 (4-7) | 26.7 (24-28.5) |

All data are mean (interquartile range, Q1-Q3). LH+ denoted days since the pre-ovulatory luteinizing hormone (LH) surge; BMI, body mass index.

**Table S4.** Demographic and clinical information for the patient cohort for Figure 7E and Figure S10C.

| Group | n = | Age | LH+ (days) | Miscarriages | BMI |
| --- | --- | --- | --- | --- | --- |
| Total | 10 | 34.6 (31.5-38.8) | 8.9 (7.8-11) | 2.7 (1-4) | 22.3 (20.1-23.8) |
| Control | 5 | 38.2 (34.5-42) | 8 (6.5-9.5) | 1.8 (0.5-4) | 23 (22-24) |
| RMM | 5 | 31 (27-34) | 9.8 (8.5-11) | 3.6 (3-4) | 21.9 (18.9-25.5) |

All data are mean (interquartile range, Q1-Q3). LH+ denoted days since the pre-ovulatory luteinizing hormone (LH) surge; BMI, body mass index.

**Table S5.**
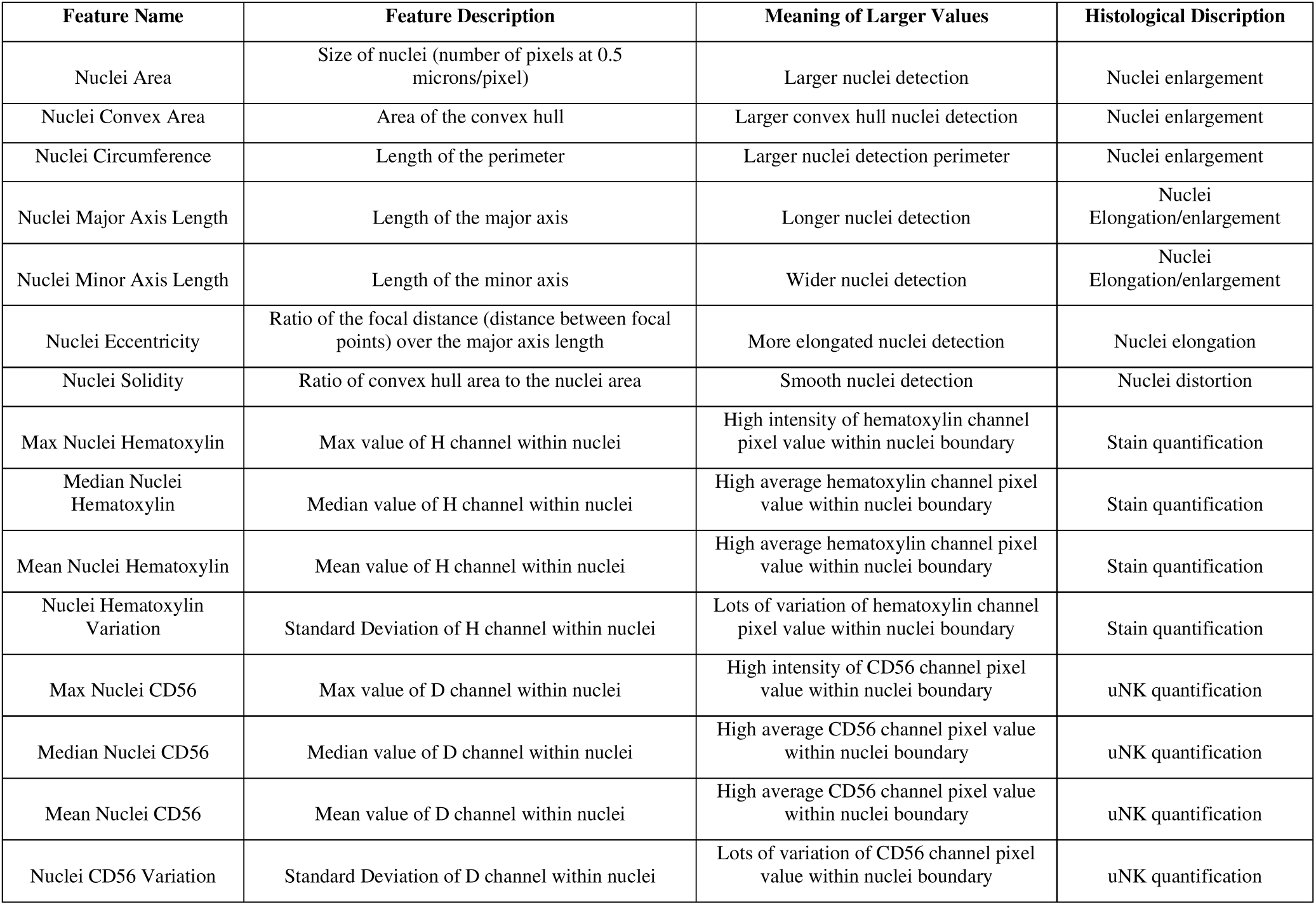
Nuclei feature descriptions and corresponding histological interpretations.

**Table S6.**
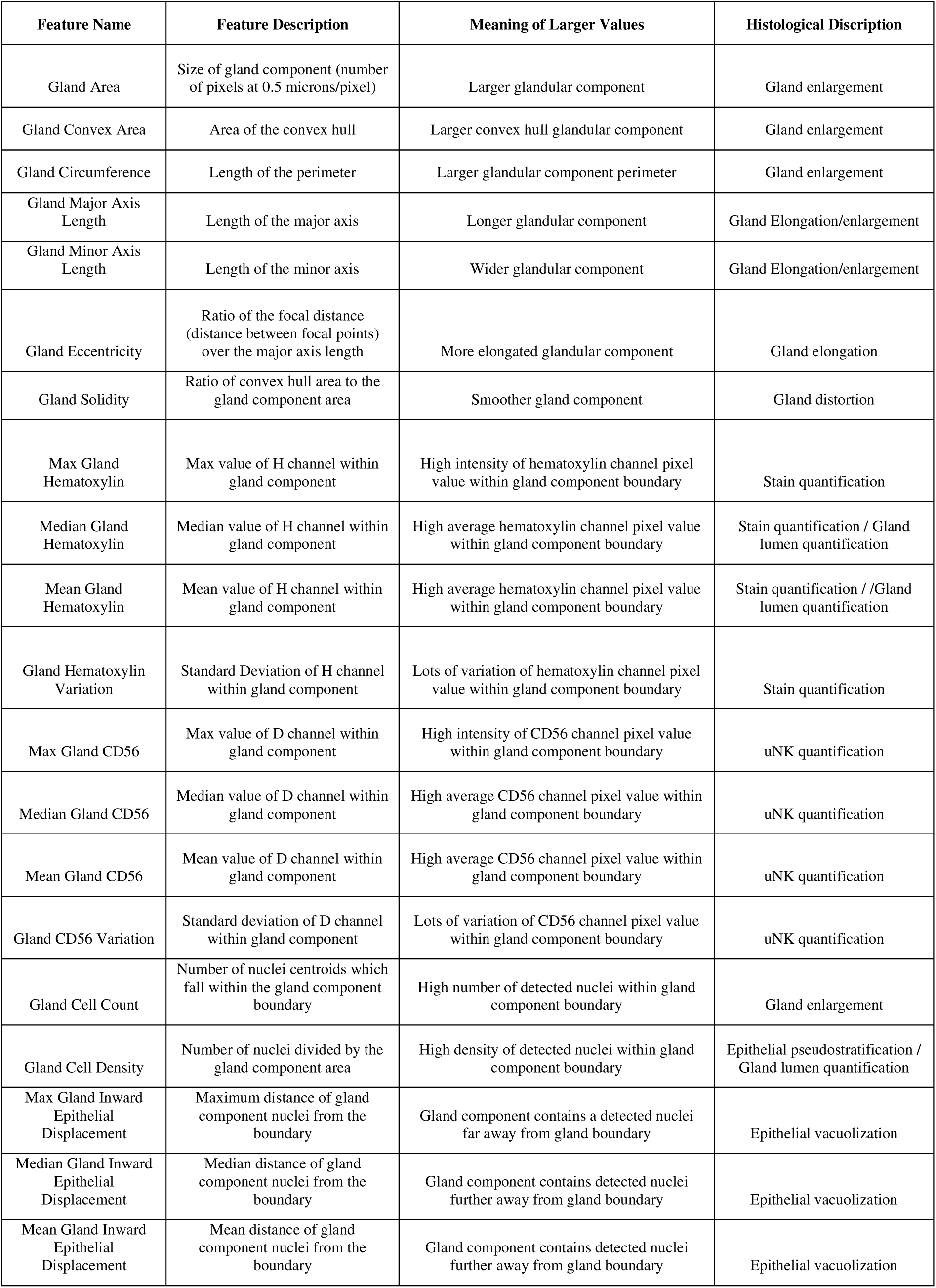
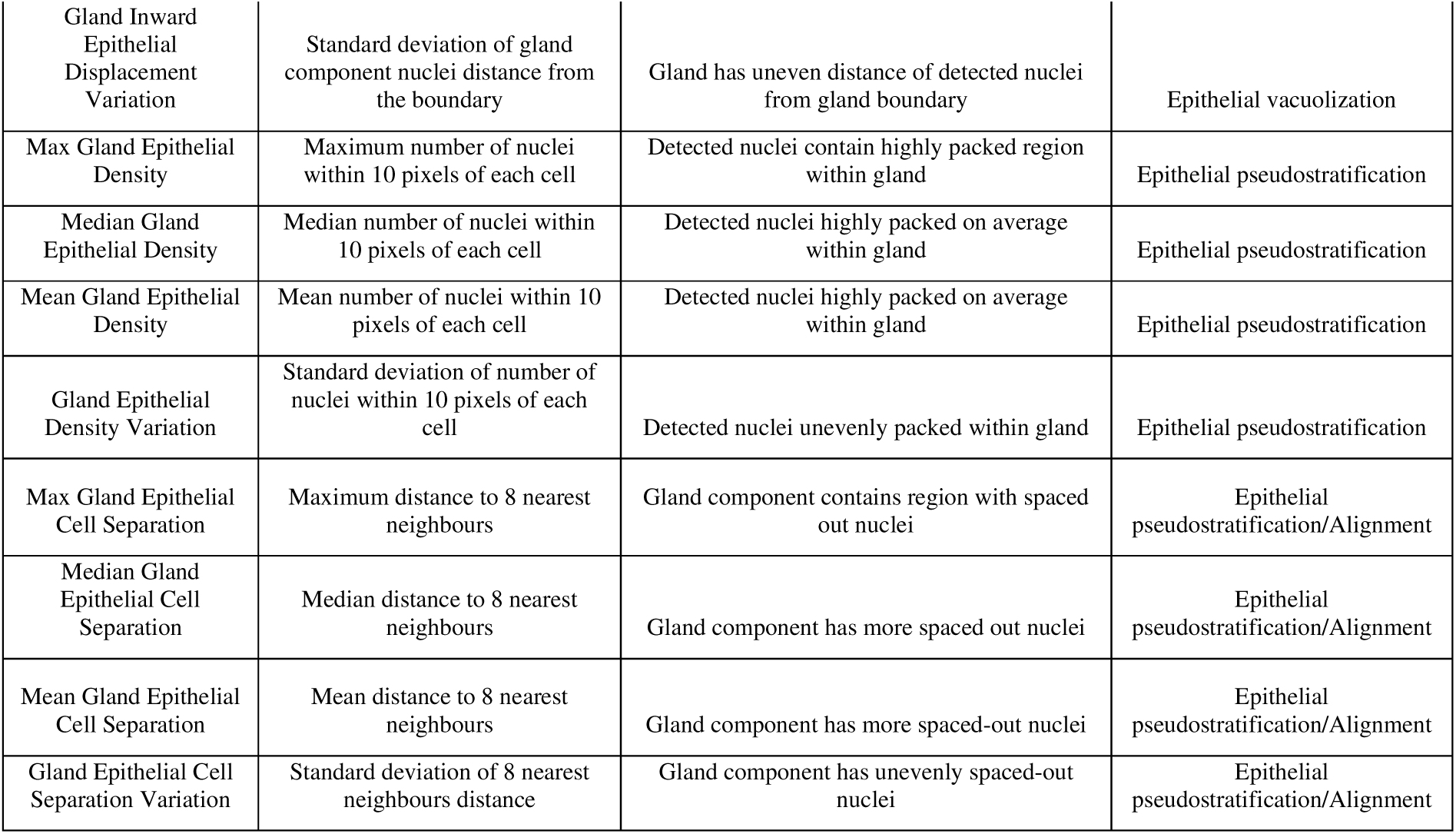
Gland component feature descriptions and corresponding histological interpretations.

### SUPPLEMENTARY FIGURES

**Figure S1.**
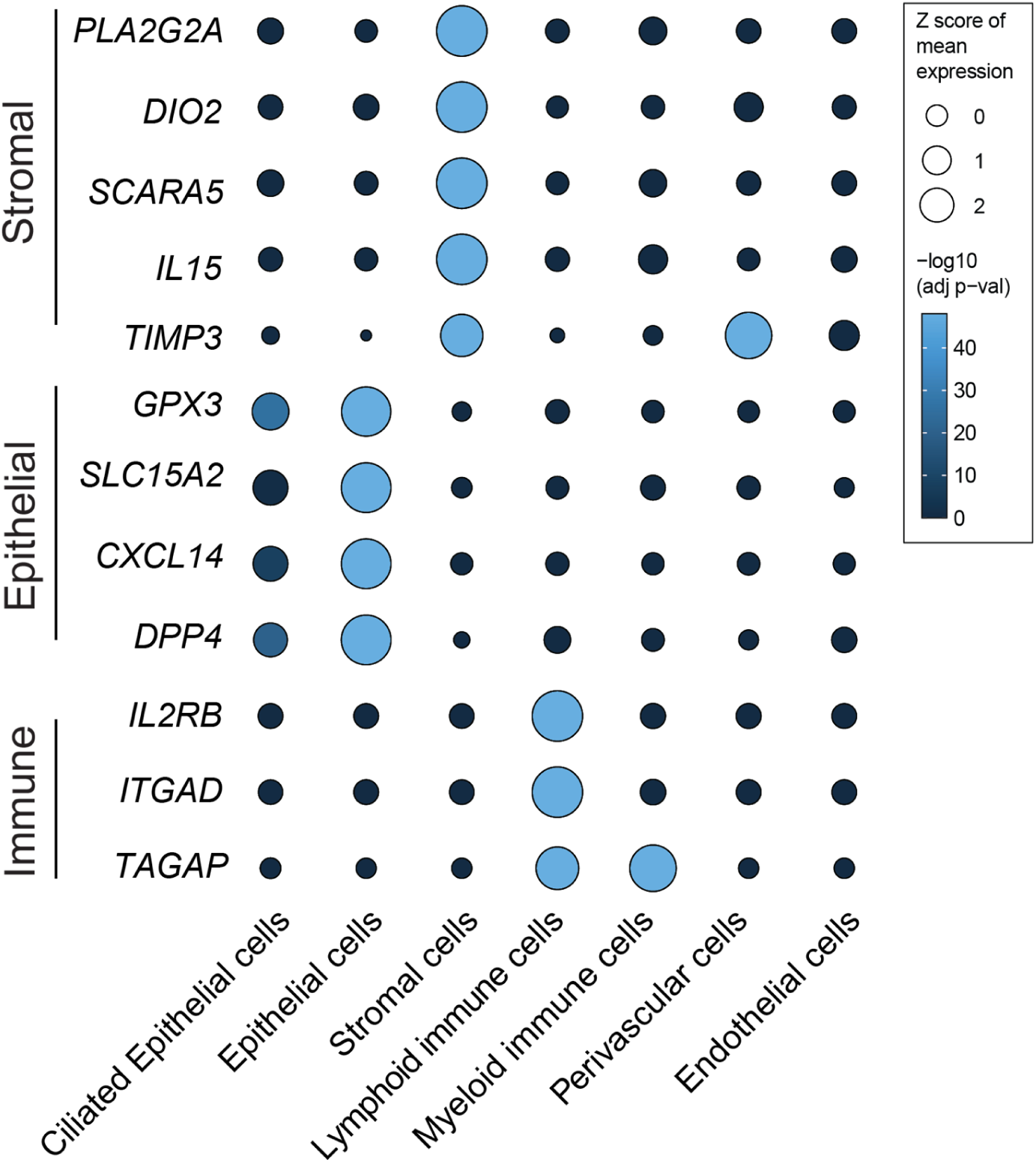
Cell-type enrichment of marker genes. Dot plot showing relative expression levels of marker genes in different midluteal endometrial cell constituents. Dot size represents z-score of the means for each gene. The colour key represents −log10 Bonferroni-adjusted Wilcoxon Rank Sum *p*-value. The data were derived from NCBI Gene Expression Omnibus repository (www.ncbi.nlm.nih.gov); accession number GSE247962.

**Figure S2.**
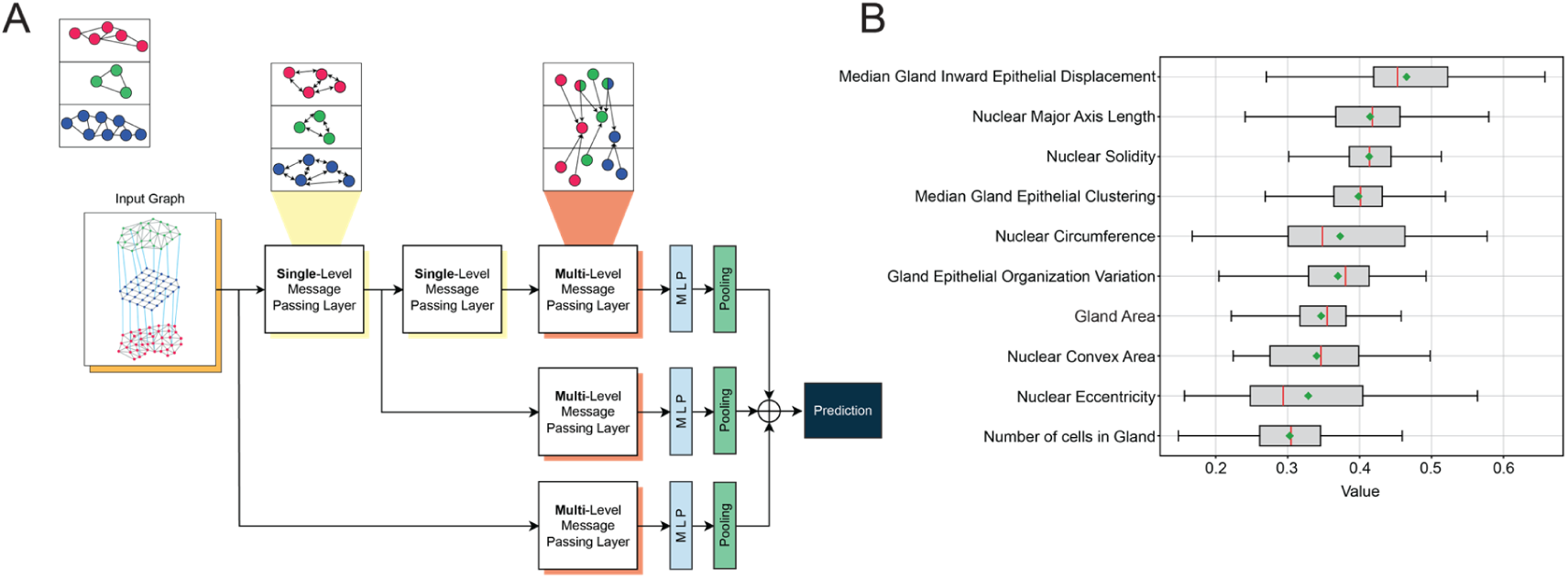
Machine learning model overview and explainability. (A) Overview of the heterogeneous graph neural network (HGNN) architecture used to predict timing and gene expression from histology images. The model integrates multi-scale patch, glandular and nuclear representations through graph-based message passing. (B) GNNExplainer output summarized as a boxplot of feature importance scores, identifying the most influential glandular and nuclear features contributing to the model output. Gland displacement from the boundary shows the highest importance value along with nuclear solidity and major axis length.

**Figure S3.**
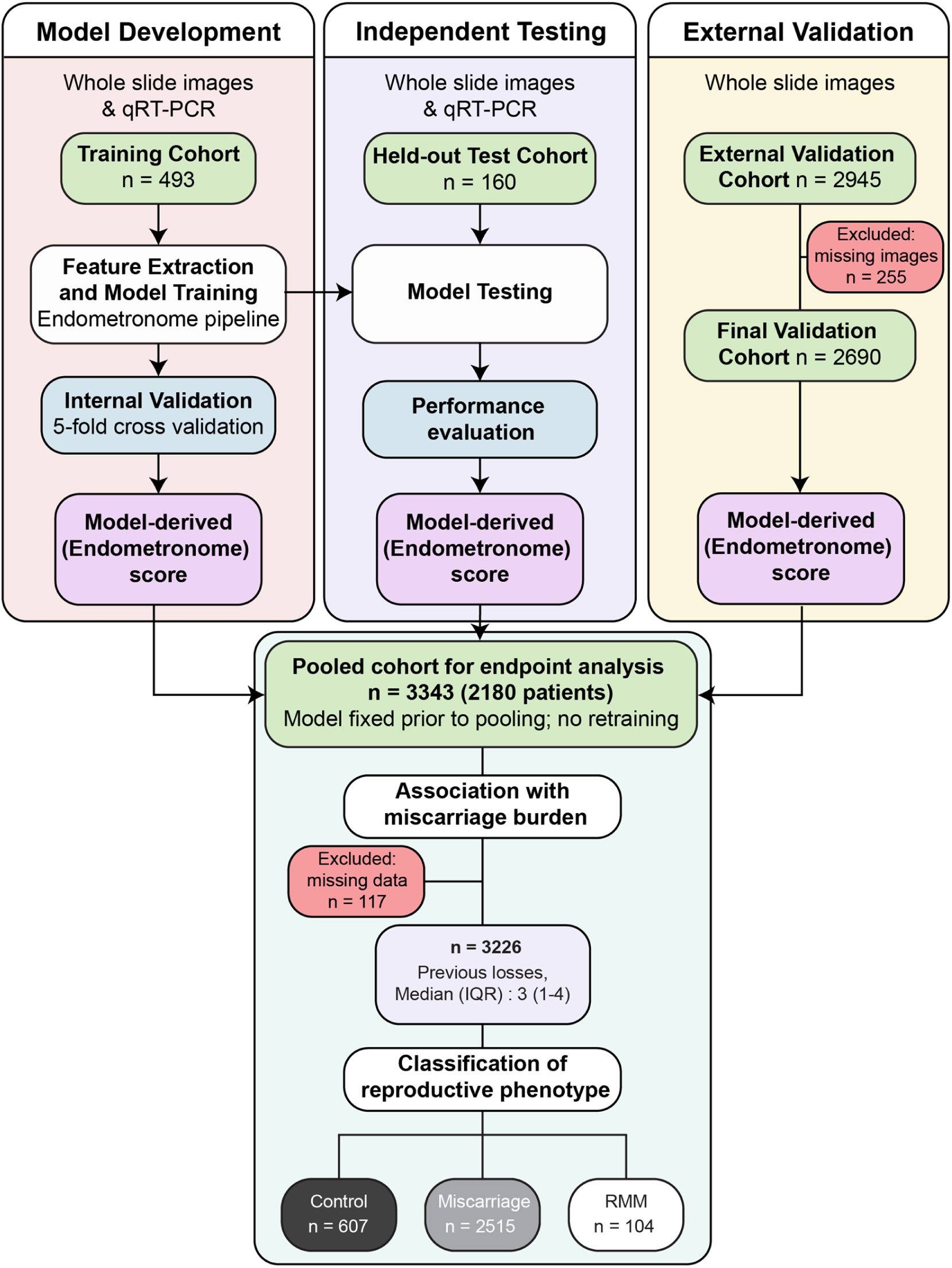
CONSORT diagram of sample inclusion and analysis workflow. Flow diagram outlining sample collection, exclusion criteria, and analysis workflow, including the allocation of samples for histology and RT–qPCR, and the final numbers used for model training, validation, and downstream analyses.

**Figure S4.**
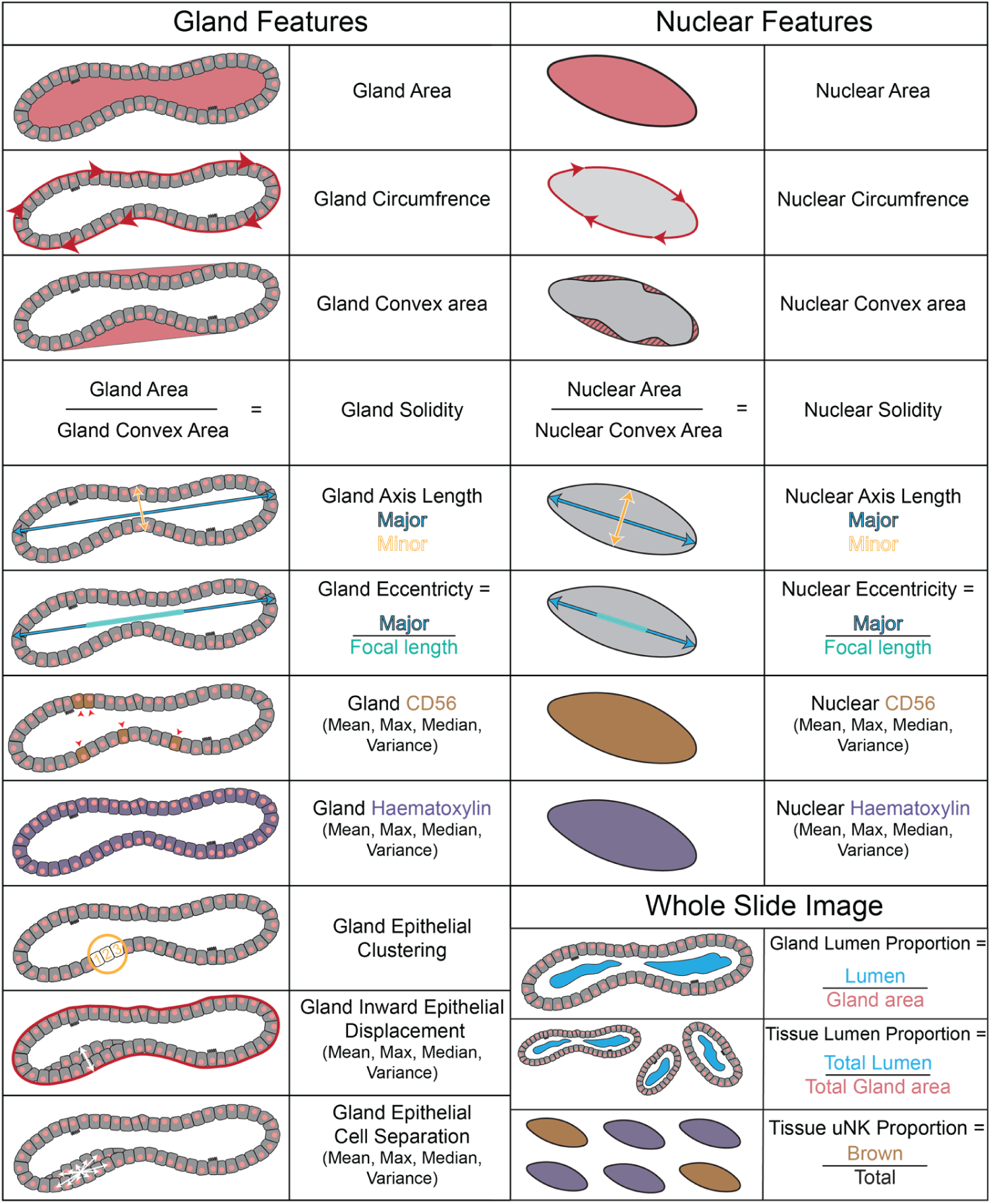
Overview illustrating each feature and its morphological interpretation for interpretability of the model and analysis.

**Figure S5.**
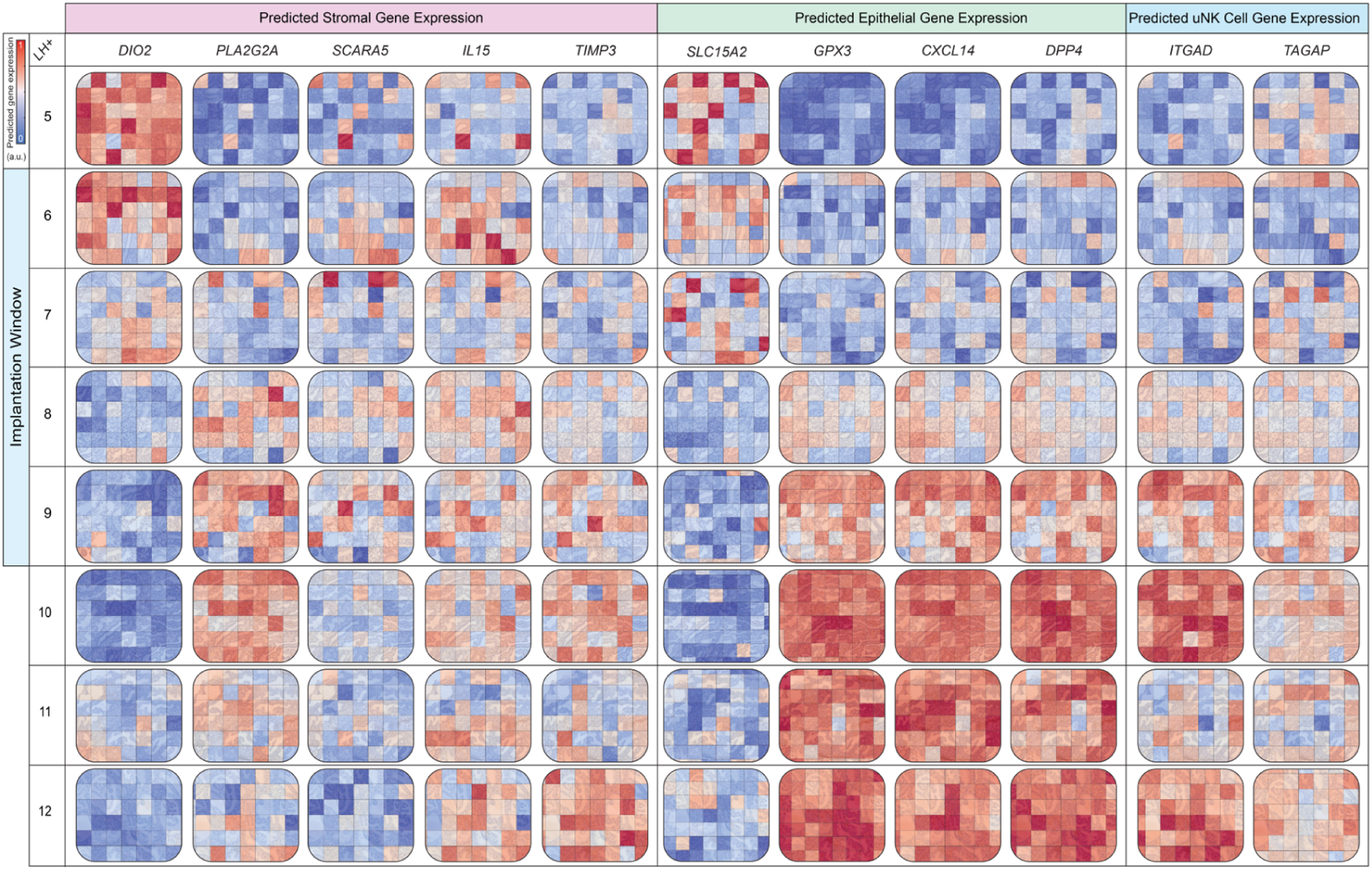
Example patch predictions of marker gene expression. Example image patches from WSIs illustrating model-predicted expression levels across the luteal phase for genes other than *GPX3* and *SLC15A2*. Representative patches are shown for various post–luteinizing hormone (LH) surge time points. Predicted expression intensities are colour-coded from low (blue) to high (red), demonstrating consistent temporal progression patterns across multiple gene targets.

**Figure S6.**
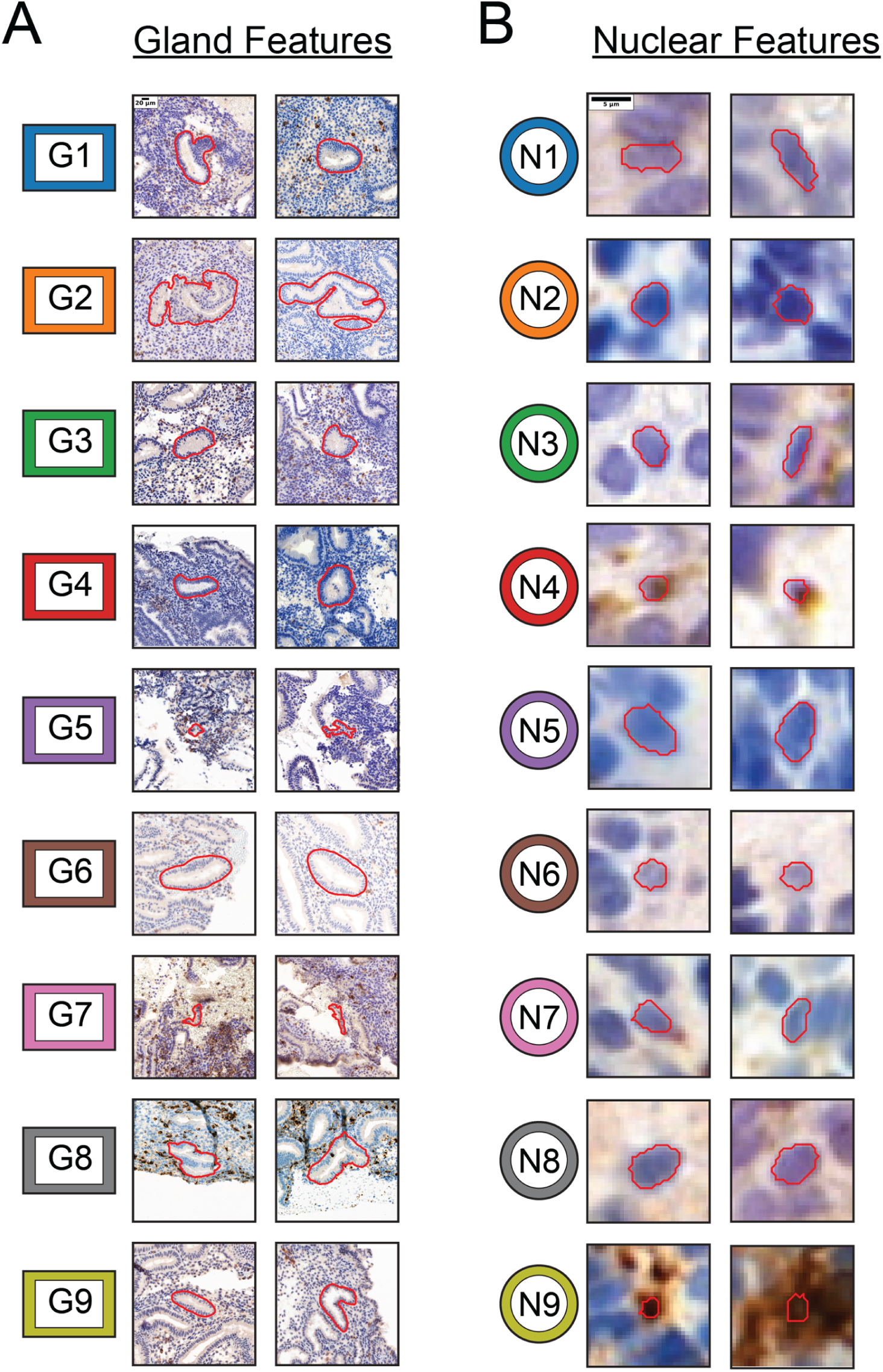
Examples of medoid gland and nuclei from each cluster. These examples highlights the interpretability of identified clusters. Cluster G3 shows very small gland components whereas cluster N9 shows high CD56 staining.

**Figure S7.**
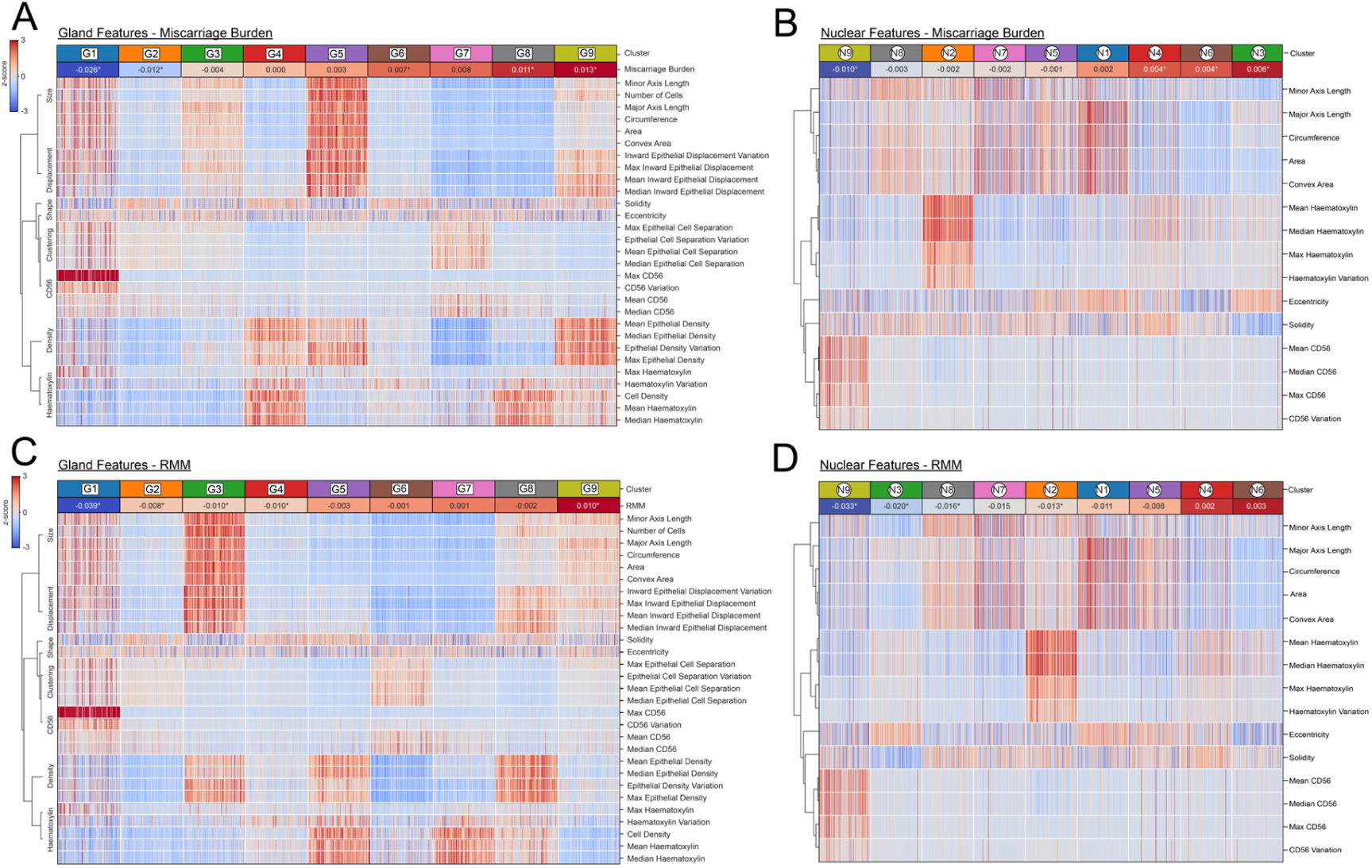
Heatmaps of glandular and nuclear feature associations with miscarriage and recurrent missed miscarriage. Z-score–normalized features are shown as heatmaps, with rows (features) hierarchically clustered by similarity and columns (clusters from k-means) ordered by regression coefficients for (A) gland clusters ordered by miscarriage burden, (B) nuclei clusters ordered by miscarriage burden, (C) gland clusters ordered by recurrent missed miscarriage (RMM), and (D) nuclei clusters ordered by RMM. The four panels illustrate distinct combinations of glandular and nuclear features changes in miscarriage and RMM outcomes, revealing differential cluster-associated patterns linked to each condition.

**Figure S8.**
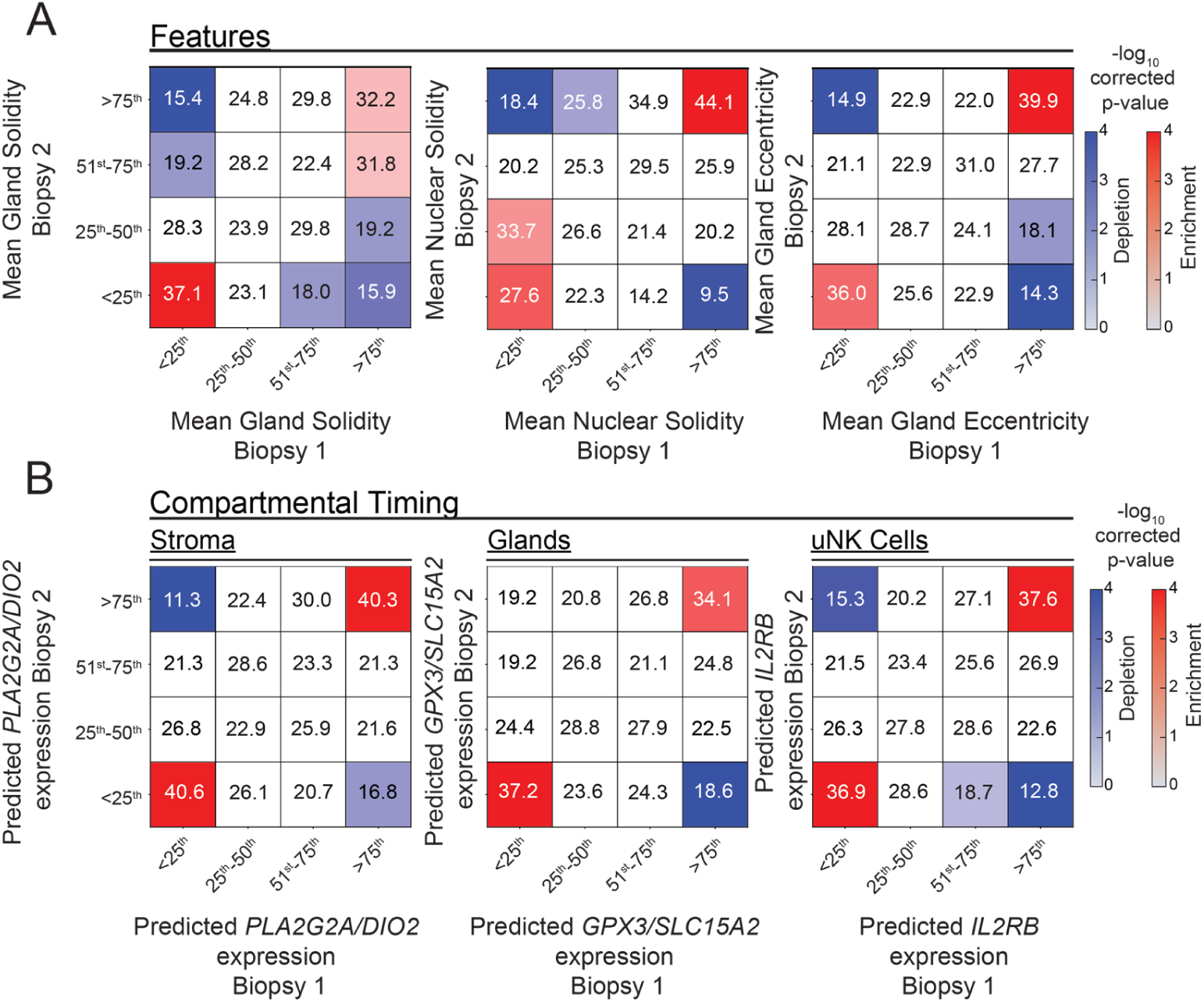
Contingency tables showing recurrence rates in paired biopsies. The coloured squares in the contingency tables indicate statistical significance (*p* < 0.05), as determined by Fisher’s exact test for enriched (red key) and depleted (blue key) associations for (A) features and (B) predicted compartmental timing. This shows that the extracted features and the subsequent model predictions are persistent over cycles.

**Figure S9.**
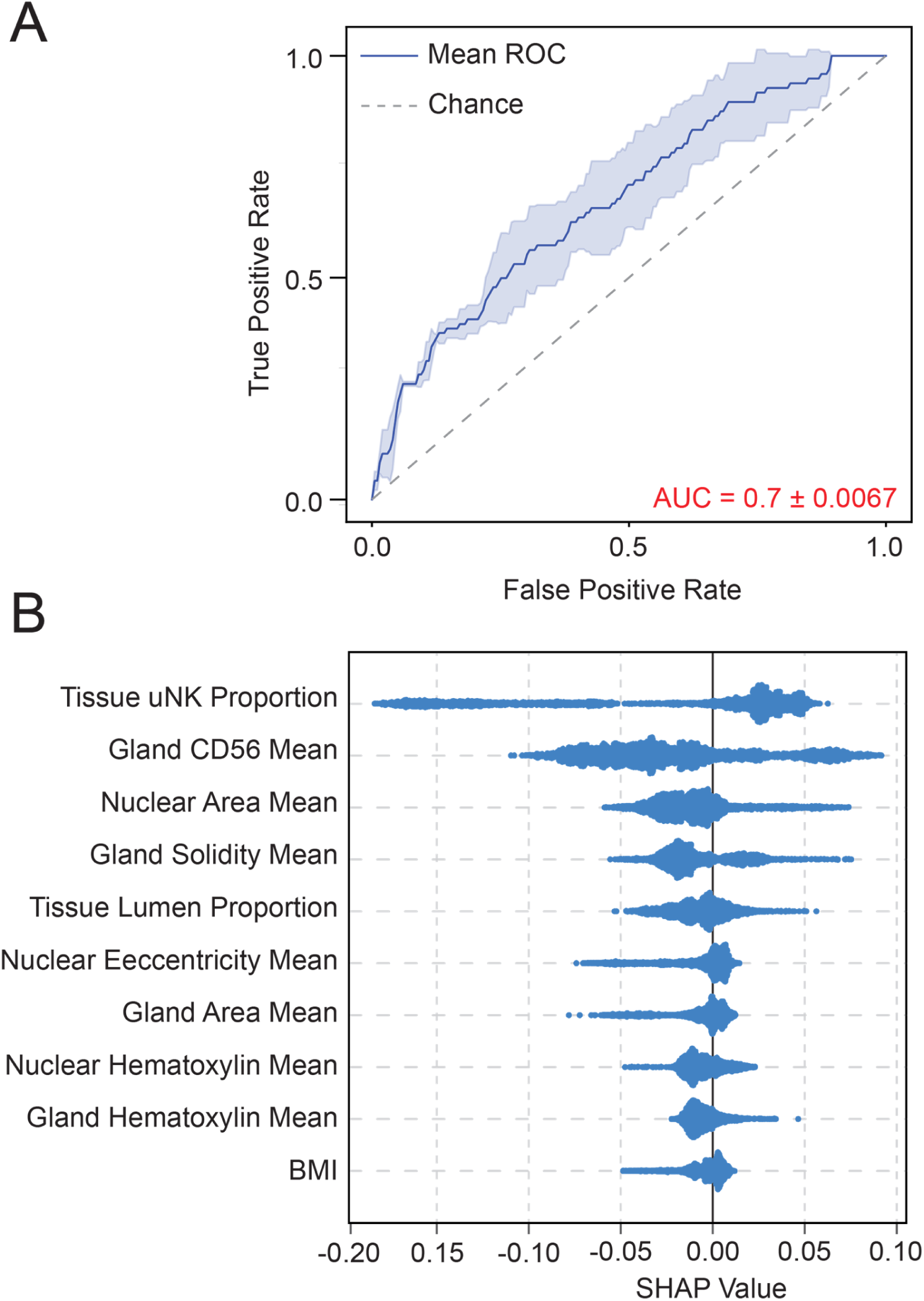
ROC curve and feature importance for predicting recurrent missed miscarriage. (A) ROC curve for random forest model predicting recurrent missed miscarriage (RMM) from image features showing the ability of predicting clincal phenotypes directly from interpretable histology image features. (B) SHAP value summary plot for the random forest model. Each point represents a sample’s contribution to model output for a given feature. Features with higher absolute SHAP values exert greater influence on model predictions. Tissue uNK cell proportion and gland CD56 staining show the highest importance values.

**Figure S10.**
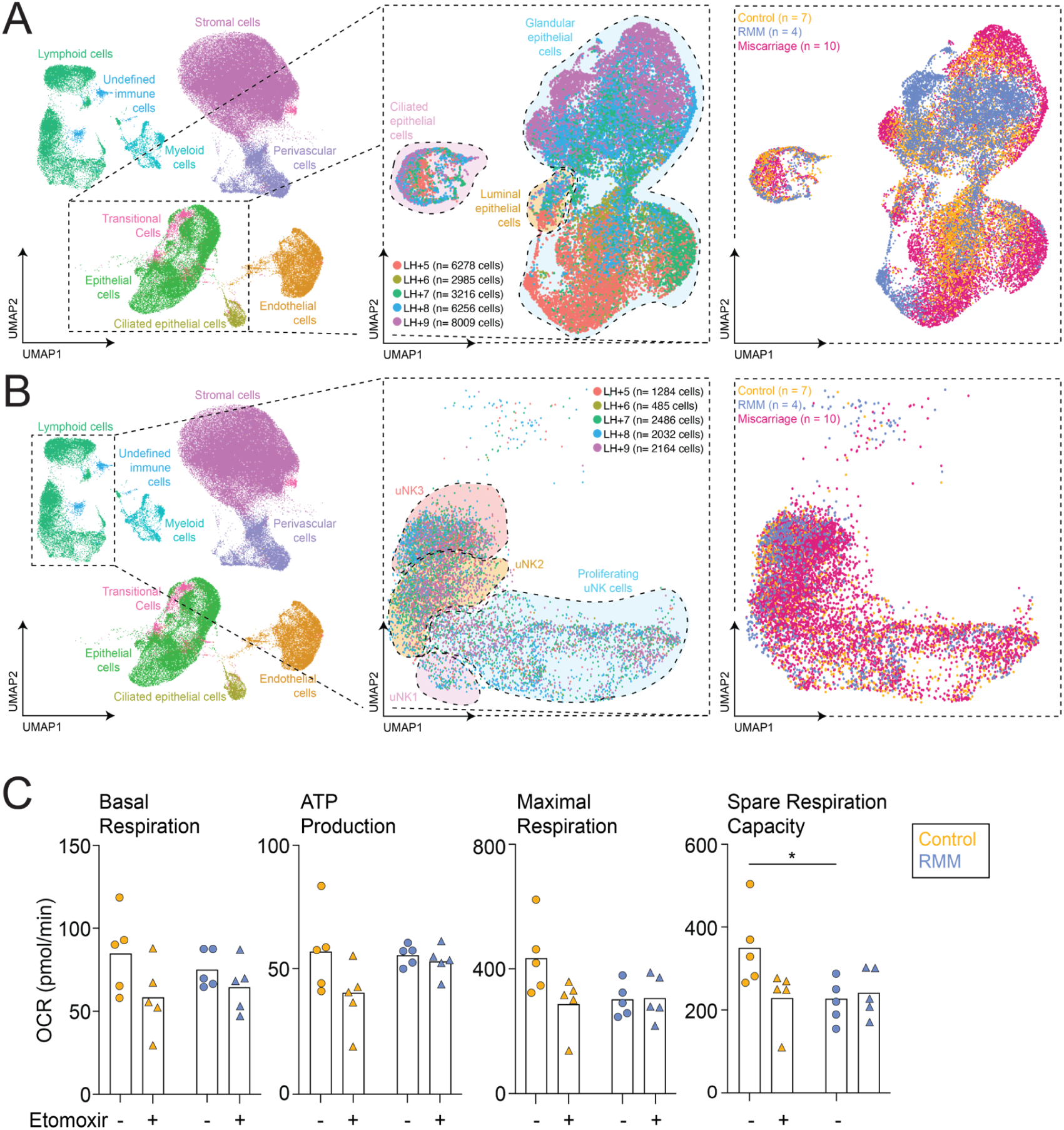
Single-cell mapping and mitochondrial dysfunction of epithelial and uNK populations in RMM endometrium. (A) Uniform manifold approximation and projection (UMAP) of scRNA-seq data from luteal phase endometrium, showing the full cellular composition (left panel) and epithelial cell populations only (middle and right panels). Colour keys depict days post-LH surge (middle panel) and patient phenotype (right panel). (B) Uniform manifold approximation and projection (UMAP) of scRNA-seq data from luteal phase endometrium, showing the full cellular composition (left panel) and uNK cell populations only (middle and right panels). Colour keys depict days post-LH surge (middle panel) and patient phenotype (right panel). (C) Bar charts showing mitochondrial metabolic parameters (basal respiration, ATP production, maximal respiration, and spare respiratory capacity) in control and RMM epithelial gland-like organoids. Bars show mean. * indicates *p* < 0.05 (Multiple Mann-Whitney tests).

## REFERENCES

1. Wagner, G.P., Kin, K., Muglia, L., and Pavlicev, M. (2014). Evolution of mammalian pregnancy and the origin of the decidual stromal cell. Int J Dev Biol 58, 117–126. 10.1387/ijdb.130335gw.

2. Erkenbrack, E.M., Maziarz, J.D., Griffith, O.W., Liang, C., Chavan, A.R., Nnamani, M.C., and Wagner, G.P. (2018). The mammalian decidual cell evolved from a cellular stress response. PLoS Biol 16, e2005594. 10.1371/journal.pbio.2005594.

3. Muter, J., Lynch, V.J., McCoy, R.C., and Brosens, J.J. (2023). Human embryo implantation. Development 150, dev201507. 10.1242/dev.201507.

4. Emera, D., Romero, R., and Wagner, G. (2012). The evolution of menstruation: a new model for genetic assimilation: explaining molecular origins of maternal responses to fetal invasiveness. Bioessays 34, 26–35. 10.1002/bies.201100099.

5. Martin, R.D. (2007). The evolution of human reproduction: a primatological perspective. Am J Phys Anthropol Suppl 45, 59–84. 10.1002/ajpa.20734.

6. Catalini, L., and Fedder, J. (2020). Characteristics of the endometrium in menstruating species: lessons learned from the animal kingdom†. Biol Reprod 102, 1160–1169. 10.1093/biolre/ioaa029.

7. Gellersen, B., and Brosens, J.J. (2014). Cyclic decidualization of the human endometrium in reproductive health and failure. Endocrine reviews 35, 851–905.

8. Salamonsen, L.A., Hutchison, J.C., and Gargett, C.E. (2021). Cyclical endometrial repair and regeneration. Development 148, dev199577. 10.1242/dev.199577.

9. Boyd, J.D., and Hamilton, W.J. (1970). The human placenta. Heffer Cambridge, 365 pp.

10. Raine-Fenning, N.J., Campbell, B.K., Clewes, J.S., Kendall, N.R., and Johnson, I.R. (2004). Defining endometrial growth during the menstrual cycle with three-dimensional ultrasound. BJOG 111, 944–949. 10.1111/j.1471-0528.2004.00214.x.

11. Muter, J., Kong, C.-S., Nebot, M.T., Tryfonos, M., Vrljicak, P., Brighton, P.J., Dimakou, D.B., Vickers, M., Yoshihara, H., Ott, S., et al. (2025). Stalling of the endometrial decidual reaction determines the recurrence risk of miscarriage. Sci Adv 11, eadv1988. 10.1126/sciadv.adv1988.

12. Marečková, M., Garcia-Alonso, L., Moullet, M., Lorenzi, V., Petryszak, R., Sancho-Serra, C., Oszlanczi, A., Icoresi Mazzeo, C., Wong, F.C.K., Kelava, I., et al. (2024). An integrated single-cell reference atlas of the human endometrium. Nat Genet 56, 1925–1937. 10.1038/s41588-024-01873-w.

13. Wang, W., Vilella, F., Alama, P., Moreno, I., Mignardi, M., Isakova, A., Pan, W., Simon, C., and Quake, S.R. (2020). Single-cell transcriptomic atlas of the human endometrium during the menstrual cycle. Nat Med 26, 1644–1653. 10.1038/s41591-020-1040-z.

14. Noyes, R.W., Hertig, A.T., and Rock, J. (2019). Reprint of: Dating the Endometrial Biopsy. Fertil Steril 112, e93–e115. 10.1016/j.fertnstert.2019.08.079.

15. Strunz, B., Bister, J., Jönsson, H., Filipovic, I., Crona-Guterstam, Y., Kvedaraite, E., Sleiers, N., Dumitrescu, B., Brännström, M., Lentini, A., et al. (2021). Continuous human uterine NK cell differentiation in response to endometrial regeneration and pregnancy. Sci Immunol 6, eabb7800. 10.1126/sciimmunol.abb7800.

16. Vrljicak, P., Lucas, E.S., Tryfonos, M., Muter, J., Ott, S., and Brosens, J.J. (2023). Dynamic chromatin remodeling in cycling human endometrium at single-cell level. Cell Rep 42, 113525. 10.1016/j.celrep.2023.113525.

17. Hertig, A.T., Rock, J., and Adams, E.C. (1956). A description of 34 human ova within the first 17 days of development. Am J Anat 98, 435–493. 10.1002/aja.1000980306.

18. Moser, G., Windsperger, K., Pollheimer, J., de Sousa Lopes, S.C., and Huppertz, B. (2018). Human trophoblast invasion: new and unexpected routes and functions. Histochem Cell Biol 150, 361–370. 10.1007/s00418-018-1699-0.

19. Burton, G.J., Watson, A.L., Hempstock, J., Skepper, J.N., and Jauniaux, E. (2002). Uterine glands provide histiotrophic nutrition for the human fetus during the first trimester of pregnancy. J Clin Endocrinol Metab 87, 2954–2959. 10.1210/jcem.87.6.8563.

20. Burton, G.J., Cindrova-Davies, T., and Turco, M.Y. (2020). Review: Histotrophic nutrition and the placental-endometrial dialogue during human early pregnancy. Placenta 102, 21–26. 10.1016/j.placenta.2020.02.008.

21. Marcy, M. (1986). Two short-term toxicity tests for the calanoid copepod Eurytemora herdmani using a complex effluent. Arch Environ Contam Toxicol 15, 199–205. 10.1007/BF01059968.

22. Pijnenborg, R., Dixon, G., Robertson, W.B., and Brosens, I. (1980). Trophoblastic invasion of human decidua from 8 to 18 weeks of pregnancy. Placenta 1, 3–19. 10.1016/s0143-4004(80)80012-9.

23. Brosens, I., Pijnenborg, R., Vercruysse, L., and Romero, R. (2011). The “Great Obstetrical Syndromes” are associated with disorders of deep placentation. Am J Obstet Gynecol 204, 193–201. 10.1016/j.ajog.2010.08.009.

24. Cao, D., Liu, Y., Cheng, Y., Wang, J., Zhang, B., Zhai, Y., Zhu, K., Liu, Y., Shang, Y., Xiao, X., et al. (2025). Time-series single-cell transcriptomic profiling of luteal-phase endometrium uncovers dynamic characteristics and its dysregulation in recurrent implantation failures. Nat Commun 16, 137. 10.1038/s41467-024-55419-z.

25. Berkhout, R.P., Lambalk, C.B., Repping, S., Hamer, G., and Mastenbroek, S. (2020). Premature expression of the decidualization marker prolactin is associated with repeated implantation failure. Gynecol Endocrinol 36, 360–364. 10.1080/09513590.2019.1650344.

26. Magnus, M.C., Wilcox, A.J., Morken, N.-H., Weinberg, C.R., and Håberg, S.E. (2019). Role of maternal age and pregnancy history in risk of miscarriage: prospective register based study. BMJ 364, l869. 10.1136/bmj.l869.

27. Quenby, S., Gallos, I.D., Dhillon-Smith, R.K., Podesek, M., Stephenson, M.D., Fisher, J., Brosens, J.J., Brewin, J., Ramhorst, R., Lucas, E.S., et al. (2021). Miscarriage matters: the epidemiological, physical, psychological, and economic costs of early pregnancy loss. The Lancet 397, 1658–1667. 10.1016/S0140-6736(21)00682-6.

28. Kolte, A.M., Westergaard, D., Lidegaard, Ø., Brunak, S., and Nielsen, H.S. (2021). Chance of live birth: a nationwide, registry-based cohort study. Hum Reprod 36, 1065–1073. 10.1093/humrep/deaa326.

29. Risch, H.A., Weiss, N.S., Clarke, E.A., and Miller, A.B. (1988). Risk factors for spontaneous abortion and its recurrence. Am J Epidemiol 128, 420–430. 10.1093/oxfordjournals.aje.a114982.

30. Garrido-Gómez, T., Castillo-Marco, N., Cordero, T., and Simón, C. (2022). Decidualization resistance in the origin of preeclampsia. Am J Obstet Gynecol 226, S886–S894. 10.1016/j.ajog.2020.09.039.

31. Muñoz-Blat, I., Pérez-Moraga, R., Castillo-Marco, N., Cordero, T., Ochando, A., Ortega-Sanchís, S., Parras-Moltó, M., Monfort-Ortiz, R., Satorres-Perez, E., Novillo, B., et al. (2025). Multi-omics-based mapping of decidualization resistance in patients with a history of severe preeclampsia. Nat Med 31, 502–513. 10.1038/s41591-024-03407-7.

32. Ammerdorffer, A., McDougall, A.R.A., Tuttle, A., Rushwan, S., Chinery, L., Vogel, J.P., Goldstein, M., and Gülmezoglu, A.M. (2024). The drug drought in maternal health: an ongoing predicament. Lancet Glob Health 12, e1174–e1183. 10.1016/S2214-109X(24)00144-X.

33. Couzin-Frankel, J. (2022). The pregnancy gap. Science 375, 1216–1220. 10.1126/science.adb2029.

34. Bilinski, A., and Emanuel, N. (2025). Fewer than 1% of United States clinical drug trials enroll pregnant participants. Am J Obstet Gynecol 232, e136–e139. 10.1016/j.ajog.2024.12.028.

35. Guglielmi, G. (2026). What drugs are safe during pregnancy? There’s a shocking lack of data. Nature 650, 286–288. 10.1038/d41586-026-00331-3.

36. Dawood, M., Eastwood, M., Jahanifar, M., Young, L., Ben-Hur, A., Branson, K., Jones, L., Rajpoot, N., and Minhas, F.U.A.A. (2023). Cross-linking breast tumor transcriptomic states and tissue histology. CR Med 4. 10.1016/j.xcrm.2023.101313.

37. Pizurica, M., Zheng, Y., Carrillo-Perez, F., Noor, H., Yao, W., Wohlfart, C., Vladimirova, A., Marchal, K., and Gevaert, O. (2024). Digital profiling of gene expression from histology images with linearized attention. Nat Commun 15, 9886. 10.1038/s41467-024-54182-5.

38. Xu, H., Usuyama, N., Bagga, J., Zhang, S., Rao, R., Naumann, T., Wong, C., Gero, Z., González, J., Gu, Y., et al. (2024). A whole-slide foundation model for digital pathology from real-world data. Nature 630, 181–188. 10.1038/s41586-024-07441-w.

39. Ding, T., Wagner, S.J., Song, A.H., Chen, R.J., Lu, M.Y., Zhang, A., Vaidya, A.J., Jaume, G., Shaban, M., Kim, A., et al. (2025). A multimodal whole-slide foundation model for pathology. Nat Med 31, 3749–3761. 10.1038/s41591-025-03982-3.

40. Ma, Y., Jamdade, S., Konduri, L., and Sailem, H. (2025). AI in Histopathology Explorer for comprehensive analysis of the evolving AI landscape in histopathology. npj Digit. Med. 8, 156. 10.1038/s41746-025-01524-2.

41. Lipecki, J., Mitchell, A.E., Muter, J., Lucas, E.S., Makwana, K., Fishwick, K., Odendaal, J., Hawkes, A., Vrljicak, P., Brosens, J.J., et al. (2022). EndoTime: non-categorical timing estimates for luteal endometrium. Hum Reprod 37, 747–761. 10.1093/humrep/deac006.

42. Russell, P., Hey-Cunningham, A., Berbic, M., Tremellen, K., Sacks, G., Gee, A., and Cheerala, B. (2014). Asynchronous glands in the endometrium of women with recurrent reproductive failure. Pathology 46, 325–332. 10.1097/PAT.0000000000000111.

43. Moore, L., Leongamornlert, D., Coorens, T.H.H., Sanders, M.A., Ellis, P., Dentro, S.C., Dawson, K.J., Butler, T., Rahbari, R., Mitchell, T.J., et al. (2020). The mutational landscape of normal human endometrial epithelium. Nature 580, 640–646. 10.1038/s41586-020-2214-z.

44. Lucas, E.S., Dyer, N.P., Murakami, K., Lee, Y.H., Chan, Y.-W., Grimaldi, G., Muter, J., Brighton, P.J., Moore, J.D., Patel, G., et al. (2016). Loss of Endometrial Plasticity in Recurrent Pregnancy Loss. Stem Cells 34, 346–356. 10.1002/stem.2222.

45. Lucas, E.S., Vrljicak, P., Muter, J., Diniz-da-Costa, M.M., Brighton, P.J., Kong, C.-S., Lipecki, J., Fishwick, K.J., Odendaal, J., Ewington, L.J., et al. (2020). Recurrent pregnancy loss is associated with a pro-senescent decidual response during the peri-implantation window. Commun Biol 3, 37. 10.1038/s42003-020-0763-1.

46. Brighton, P.J., Maruyama, Y., Fishwick, K., Vrljicak, P., Tewary, S., Fujihara, R., Muter, J., Lucas, E.S., Yamada, T., Woods, L., et al. (2017). Clearance of senescent decidual cells by uterine natural killer cells in cycling human endometrium. Elife 6, e31274. 10.7554/eLife.31274.

47. Browaeys, R., Gilis, J., Sang-Aram, C., De Bleser, P., Hoste, L., Tavernier, S., Lambrechts, D., Seurinck, R., and Saeys, Y. (2023). MultiNicheNet: a flexible framework for differential cell-cell communication analysis from multi-sample multi-condition single-cell transcriptomics data (BioRxiv) 10.1101/2023.06.13.544751.

48. De Rosa, V., Iommelli, F., Monti, M., Fonti, R., Votta, G., Stoppelli, M.P., and Del Vecchio, S. (2015). Reversal of Warburg Effect and Reactivation of Oxidative Phosphorylation by Differential Inhibition of EGFR Signaling Pathways in Non-Small Cell Lung Cancer. Clin Cancer Res 21, 5110–5120. 10.1158/1078-0432.CCR-15-0375.

49. Burns, J.S., and Manda, G. (2017). Metabolic Pathways of the Warburg Effect in Health and Disease: Perspectives of Choice, Chain or Chance. Int J Mol Sci 18, 2755. 10.3390/ijms18122755.

50. Garratt, M., Gaillard, J.-M., Brooks, R.C., and Lemaître, J.-F. (2013). Diversification of the eutherian placenta is associated with changes in the pace of life. Proc Natl Acad Sci U S A 110, 7760–7765. 10.1073/pnas.1305018110.

51. Laundon, D., Gostling, N.J., Sengers, B.G., Chavatte-Palmer, P., and Lewis, R.M. (2024). Placental evolution from a three-dimensional and multiscale structural perspective. Evolution 78, 13–25. 10.1093/evolut/qpad209.

52. Afzal, J., Suhail, Y., Du, W., Liu, Y., Ramasamy, R., Liu, Z., Goyal, R., Novin, A., Suhail, S., Maziarz, J., et al. (2025). Evidence for coopetition at the maternal-fetal interface shaping placental invasion. Proc Natl Acad Sci U S A 122, e2323038122. 10.1073/pnas.2323038122.

53. Vanneste, E., Voet, T., Le Caignec, C., Ampe, M., Konings, P., Melotte, C., Debrock, S., Amyere, M., Vikkula, M., Schuit, F., et al. (2009). Chromosome instability is common in human cleavage-stage embryos. Nat Med 15, 577–583. 10.1038/nm.1924.

54. McCoy, R.C. (2017). Mosaicism in Preimplantation Human Embryos: When Chromosomal Abnormalities Are the Norm. Trends Genet 33, 448–463. 10.1016/j.tig.2017.04.001.

55. Macklon, N.S., and Brosens, J.J. (2014). The human endometrium as a sensor of embryo quality. Biol Reprod 91, 98. 10.1095/biolreprod.114.122846.

56. Evers, J.L.H. (2002). Female subfertility. Lancet 360, 151–159. 10.1016/S0140-6736(02)09417-5.

57. Magnus, M.C., Morken, N.-H., Wensaas, K.-A., Wilcox, A.J., and Håberg, S.E. (2021). Risk of miscarriage in women with chronic diseases in Norway: A registry linkage study. PLoS Med 18, e1003603. 10.1371/journal.pmed.1003603.

58. Magnus, M.C., Hockey, R.L., Håberg, S.E., and Mishra, G.D. (2022). Pre-pregnancy lifestyle characteristics and risk of miscarriage: the Australian Longitudinal Study on Women’s Health. BMC Pregnancy Childbirth 22, 169. 10.1186/s12884-022-04482-9.

59. Wang, H., and Dey, S.K. (2006). Roadmap to embryo implantation: clues from mouse models. Nat Rev Genet 7, 185–199. 10.1038/nrg1808.

60. Dimitriadis, E., Menkhorst, E., Saito, S., Kutteh, W.H., and Brosens, J.J. (2020). Recurrent pregnancy loss. Nature reviews disease primers 6, 98. 10.1038/s41572-020-00228-z.

61. Watson, D.S., Krutzinna, J., Bruce, I.N., Griffiths, C.E.M., McInnes, I.B., Barnes, M.R., and Floridi, L. (2019). Clinical applications of machine learning algorithms: beyond the black box. Bmj 364. 10.1136/bmj.l886.

62. Leslie, D. (2019). Understanding artificial intelligence ethics and safety. arXiv preprint arXiv:1906.05684. 10.48550/arXiv.1906.05684.

63. Yoon, C.H., Torrance, R., and Scheinerman, N. (2022). Machine learning in medicine: should the pursuit of enhanced interpretability be abandoned? Journal of Medical Ethics 48, 581–585. 10.1136/medethics-2020-107102.

64. Dhillon-Smith, R.K., Melo, P., Devall, A.J., Smith, P.P., Al-Memar, M., Barnhart, K., Condous, G., Christiansen, O.B., Goddijn, M., Jurkovic, D., et al. (2023). A core outcome set for trials in miscarriage management and prevention: An international consensus development study. BJOG 130, 1346–1354. 10.1111/1471-0528.17484.

65. Wright, G., Brighton, P., Yoshihara, H., Thornton, J., Muter, J., Brosens, J., and Minhas, F. (2025). ASTER: Automated Segmentation of Endometrial Histology Images for Reproductive Health Assessment. In Annual Conference on Medical Image Understanding and Analysis (Springer), pp. 43–57. 10.1007/978-3-031-98688-8_4.

66. Ma, N., Zhang, X., Zheng, H.-T., and Sun, J. (2018). ShuffleNet V2: Practical Guidelines for Efficient CNN Architecture Design. CoRR abs/1807.11164. 10.48550/arXiv.1807.11164.

67. Russakovsky, O., Deng, J., Su, H., Krause, J., Satheesh, S., Ma, S., Huang, Z., Karpathy, A., Khosla, A., Bernstein, M., et al. (2015). Imagenet large scale visual recognition challenge. International journal of computer vision 115, 211–252. 10.48550/arXiv.1409.0575.

68. Wang, Y., Sun, Y., Liu, Z., Sarma, S.E., Bronstein, M.M., and Solomon, J.M. (2019). Dynamic graph cnn for learning on point clouds. Acm Transactions On Graphics (tog) 38, 1–12. 10.48550/arXiv.1801.07829.

69. Xu, K., Hu, W., Leskovec, J., and Jegelka, S. (2018). How powerful are graph neural networks? arXiv preprint arXiv:1810.00826. 10.48550/arXiv.1810.00826.

70. Kingma, D.P., and Ba, J. (2014). Adam: A method for stochastic optimization. arXiv preprint arXiv:1412.6980. 10.48550/arXiv.1412.6980.

71. Ying, Z., Bourgeois, D., You, J., Zitnik, M., and Leskovec, J. (2019). Gnnexplainer: Generating explanations for graph neural networks. Advances in neural information processing systems 32. 10.48550/arXiv.1903.03894.

72. McInnes, L., Healy, J., and Melville, J. (2018). Umap: Uniform manifold approximation and projection for dimension reduction. arXiv preprint arXiv:1802.03426. 10.48550/arXiv.1802.03426.

73. Wright, G., Keller, P., Muter, J., Brosens, J.J., Tejpar, S., and Minhas, F. (2026). DCAFA: Differential Community Abundance and Feature Analysis for Histological Images. bioRxiv 2026.04.28.721329. 10.64898/2026.04.28.721329.

74. Lundberg, S.M., and Lee, S.-I. (2017). A Unified Approach to Interpreting Model Predictions. Advances in neural information processing systems 30. 10.48550/arXiv.1705.07874.

75. Turco, M.Y., Gardner, L., Hughes, J., Cindrova-Davies, T., Gomez, M.J., Farrell, L., Hollinshead, M., Marsh, S.G.E., Brosens, J.J., Critchley, H.O., et al. (2017). Long-term, hormone-responsive organoid cultures of human endometrium in a chemically defined medium. Nat Cell Biol 19, 568–577. 10.1038/ncb3516.

